# Inter-species variation in number of bristles on forewings of tiny insects does not impact clap-and-fling aerodynamics

**DOI:** 10.1101/2020.10.27.356337

**Authors:** Vishwa T. Kasoju, Mitchell P. Ford, Truc T. Ngo, Arvind Santhanakrishnan

## Abstract

Flight-capable miniature insects of body length (BL) < 2 mm typically possess wings with long bristles on the fringes. Though their flight is challenged by needing to overcome significant viscous resistance at chord-based Reynolds number (*Re_c_*) on the order of 10, these insects use clap-and-fling mechanism coupled with bristled wings for lift augmentation and drag reduction. However, inter-species variation in the number of bristles (*n*) and inter-bristle gap (*G*) to bristle diameter (*D*) ratio (*G*/*D*) and their effects on clap-and-fling aerodynamics remain unknown. Forewing image analyses of 16 species of thrips and 21 species of fairyflies showed that *n* and maximum wing span were both positively correlated with BL. We conducted aerodynamic force measurements and flow visualization on simplified physical models of bristled wing pairs that were prescribed to execute clap-and-fling kinematics at *Re*_c_=10 using a dynamically scaled robotic platform. 23 bristled wing pairs were tested to examine the isolated effects of changing dimensional (*G, D*, span) and non-dimensional (*n, G/D*) geometric variables on dimensionless lift and drag. Within biologically observed ranges of *n* and *G*/*D*, we found that: (a) increasing *G* provided more drag reduction than decreasing *D*; (b) changing *n* had minimal impact on lift generation; and (c) varying *G/D* produced minimal changes in aerodynamic forces. Taken together with the broad variation in *n* (32-161) across the species considered here, the lack of impact of changing *n* on lift generation suggests that tiny insects may experience reduced biological pressure to functionally optimize *n* for a given wing span.

**SUMMARY STATEMENT:** Integrating morphological analysis of bristled wings seen in miniature insects with physical model experiments, we find that aerodynamic forces are unaffected across the broad biological variation in number of bristles.

## INTRODUCTION

The wings of flying insects show tremendous diversity in shape, size and function. Curiously, the wings of several families of flight-capable miniature insects smaller than fruit flies have independently evolved *ptiloptery* (Polilov, 2015; Sane, 2016), resulting in wings with long setae at the fringes. Though their extremely small sizes (body length < 2 mm) make visual observation difficult, tiny flying insects are not limited to just a few outlying examples. Rather, more than 5,500 species of Thysanoptera (thrips) (Morse and Hoddle, 2006), as well as several hundred species of Mymaridae (fairyflies) and Trichogrammatidae have been identified to date. Despite their agricultural and ecological importance in acting as biological vectors of plant viruses and as invasive pests of commercially important plants (Ullman et al., 2002; Jones, 2005), our understanding of the flight biomechanics of tiny insects is far from complete. Due to the difficulty in acquiring free-flight recordings of tiny insects, several studies have used physical and computational modeling to examine the functional significance of wing bristles (Santhanakrishnan et al., 2014; Jones et al., 2016; Lee and Kim, 2017; Kasoju et al., 2018). However, little is known about the extent of variation in bristled wing morphology among different species of tiny insects. It remains unclear whether tiny insects experience biological pressure to optimize the mechanical design of their bristled wings toward improving flight aerodynamics.

Pronounced viscous dissipation of kinetic energy occurs at wing length scales on the order of 1 mm, making it difficult for tiny insects to stay aloft. The relative importance of inertial to viscous forces in a fluid flow is characterized using the dimensionless Reynolds number (*Re*):

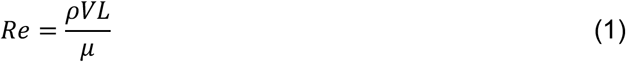

where *ρ* and *μ* are the density and dynamic viscosity of the fluid medium, respectively; *V* and *L* are characteristic velocity and length scales, respectively. Tiny insects typically operate at wing chord (*c*) based *Re* (*Re_c_* = *ρVc*/*μ*) on the orders of 1 to 10 and bristle diameter (*D*) based *Re* (*Re_b_* = *ρVD/μ*) ranging between 0.01-0.07 (Ellington, 1975; Kuethe, 1975; Santhanakrishnan et al., 2014; Jones et al., 2016). Despite the difficulty in sustaining flight at such low *Re_c_*, entomological studies have reported active flight and dispersal of thrips (Morse and Hoddle, 2006; Rodriguez-Saona et al., 2010). Tiny insects use biomechanical adaptations to overcome the fluid dynamic challenges associated with flight at small scales. These insects operate their wings at near-maximum stroke amplitude using the ‘clap-and-fling’ mechanism, first observed by Weis-Fogh (1973) in *Encarsia formosa*. The use of clap-and-fling has been documented in other freely flying tiny insects, including *Thrips physapus* (Ellington, 1975) and *Muscidifurax raptor* (Miller and Peskin, 2009). Wing rotation during fling has been noted to augment lift via the generation of a leading edge vortex (LEV) on the wings (Weis-Fogh, 1973; Lighthill, 1973; Spedding and Maxworthy, 1986; Miller and Peskin, 2005; Lehmann et al., 2005; Lehmann and Pick, 2007; Miller and Peskin, 2009; Arora et al., 2014). However, the concomitant generation of large drag force at the start of fling undermines the lucrativeness of clap-and-fling at *Re_c_* relevant to tiny insect flight (Miller and Peskin, 2005; Arora et al., 2014). Previous studies (Santhanakrishnan et al., 2014; Jones et al., 2016; Kasoju et al., 2018; Ford et al., 2019) have shown that bristled wings can reduce the force required to fling the wings apart.

Although a number of studies have examined the flow structures and aerodynamic forces generated by bristled wings in comparison with solid wings (Sunada et al., 2002; Santhanakrishnan et al., 2014; Jones et al., 2016; Lee and Kim, 2017; Lee et al., 2018; Kasoju et al., 2018), morphological variation of bristled wing design in tiny flying insects is far less documented. Jones et al. (2016) examined the inter-bristle gap (*G*), bristle diameter (*D*), and the wing area covered by bristles in the forewings of 23 species of fairyflies (Mymaridae). With decreasing body length (BL), they found that *G* and *D* decreased and area occupied by bristles increased. Ford et al. (2019) found that the ratio of solid membrane area (*A*_M_) to total wing area (*A*_T_) in the forewings of 25 species of thrips (Thysanoptera) ranged from 14% to 27%, as compared to the *A*_M_/*A*_T_ range of 11% to 88% in smaller-sized fairyflies examined by Jones et al. (2016). Using physical models that were prescribed to execute clap-and-fling kinematics, Ford et al. (2019) found that lift to drag ratios were largest for bristled wing models with *A*_M_/*A*_T_ similar to thrips forewings. Inter-species variation of *G, D*, wing span (*S*) and number of bristles (*n*), as well as their concomitant effects on clap-and-fling aerodynamics, are currently unknown.

Due to the large number of taxa of tiny insects that possess bristled wings, we expected a broad range of variation in morphological characteristics. We hypothesized that at *Re_b_* and *Re_c_* relevant to tiny insect flight, dimensionless aerodynamic forces generated by clap-and-fling would be minimally impacted by individually varying *n* and *G/D* within their biological ranges. If true, tiny flying insects may not experience biological pressure to functionally optimize the mechanical design of their bristled wings. We measured *n* and maximum wing span (*S*_max_) from published forewing images of 16 species of thrips (Thysanoptera) and 21 species of fairyflies (Mymaridae). In addition, we measured *G* and *D* from forewing images of 22 Thysanoptera species and calculated *G/D* ratios to compare to those of smaller-sized Mymaridae species that were reported by Jones et al. (2016). The thrips and fairyfly species considered here encompass BL ranging from 0.1 mm to 2 mm, making this study relevant for a broad range of tiny flying insects. Using the morphological data, we fabricated physical bristled wing models varying in *G, D, S*, and *n*. These physical models were comparatively tested using a dynamically scaled robotic platform mimicking the portion of clap-and-fling kinematics where wing-wing interaction occurs. Aerodynamic force measurements and flow field visualization were conducted to identify the functional significance of the above bristled wing design variables.

## MATERIALS AND METHODS

### Forewing morphology

We measured average BL, *A*_T_, *S*_max_, *n, G* and *D* from published forewing images of several species of thrips (Thysanoptera) and fairyflies (Mymaridae). Jones et al., (2016) measured *G* and *D* from previously published forewing images of 23 species of Mymaridae and found the *G/D* ratio to not be correlated with BL. However, the wing span, chord and *n* of Mymaridae forewings were not reported by Jones et al. (2016) and are characterized in this study (21 species). Nearly all the Mymaridae species considered by Jones et al. (2016) were of BL less than 1 mm, while the BL of the thrips species considered here range between 1-2 mm. For all species considered here, average wing chord (*C*_ave_) was calculated from the measurements of *A*_T_ and *S*_max_.

We required that each published forewing image considered for measurements of *S*_max_, *A*_T_ and *n* met the following criteria: 1) contains a scale bar; 2) consist of least one forewing zoomed out with all bristles shown; and 3) no noticeable damage to any of the forewing bristles. We used a different set of published forewing images for measurements of *G* and *D*, as we needed to substantially magnify each of these images (as compared to measurements of *S*_max_, *A*_T_ and *n*). We required that the published forewing images considered for *G* and *D* measurements had a spatial resolution of at least 6 pixels per bristle diameter, similar to the criterion used by Jones et al. (2016). As *G* and *D* measurements were used to compute non-dimensional *G/D* ratios, we did not restrict the images selected for *G* and *D* measurements to only those that contained a scale bar (as measurements of *G* and *D* in pixels from a forewing image would suffice to calculate the dimensionless *G*/*D* ratio).

Based on the above criteria, forewing images of 16 thrips species were selected for measuring *S*_max_, *A*_T_ and *n*, and of 22 thrips species for measuring *G* and *D* (Mound & Reynaud, 2005; Mound, 2009; Zang et al., 2010; Riley et al., 2011; MAF Plant Health & Environment Laboratory, 2011; Cavalleri and Mound, 2012; Ng and Mound, 2012; Masumoto & Okajima 2013; Minaei and Aleosfoor, 2013; Zamar et al., 2013; Cavalleri and Mound, 2014; Dang et al., 2014; Ng and Mound, 2015; Cavalleri et al., 2016; Lima and Mound, 2016; Mound and Tree, 2016; Wang and Tang, 2016; Goldaracene & Hance 2017). The thrips species considered here encompass three different taxonomic families. In addition, 21 Mymaridae species were selected for measuring *S*_max_, *A*_T_ and *n* (Huber, Mendel et al., 2006; Huber & Baquero, 2007; Lin et al., 2007; Huber, Gibson et al., 2008; Huber & Noyes 2013).

Bristled wing morphological variables were measured from these images using ImageJ software (Schneider et al., 2012). *S*_max_ was defined to be the distance from the center of the wing root to the tip of the bristles, and was measured using ImageJ according to the diagram in Fig. 1A. Average wing chord (*C*_ave_) was calculated by measuring *A*_T_ using the same procedure as in Jones et al. (2016) and Ford et al. (2019) and dividing *A*_T_ by *S*_max_. As the forewing images obtained from the various sources were aligned in different orientations, we rotated the wings before measurements such that they were always oriented horizontally. *G*/*D* ratio was calculated from the measurements of *G* and *D* in the forewing images. BL measurements were made either based on the scale bar (where available), or from the text of the article containing the image. The measured values were plotted against BL (*S*max and *n* in Fig. 1B,C; *G/D* in Fig. 1D). For each measured quantity, linear regressions were performed and R^2^ and *p*-values were determined. A full list of species and corresponding measurements are provided as supplementary material (Tables S1,S2,S3).

**Figure 1.**
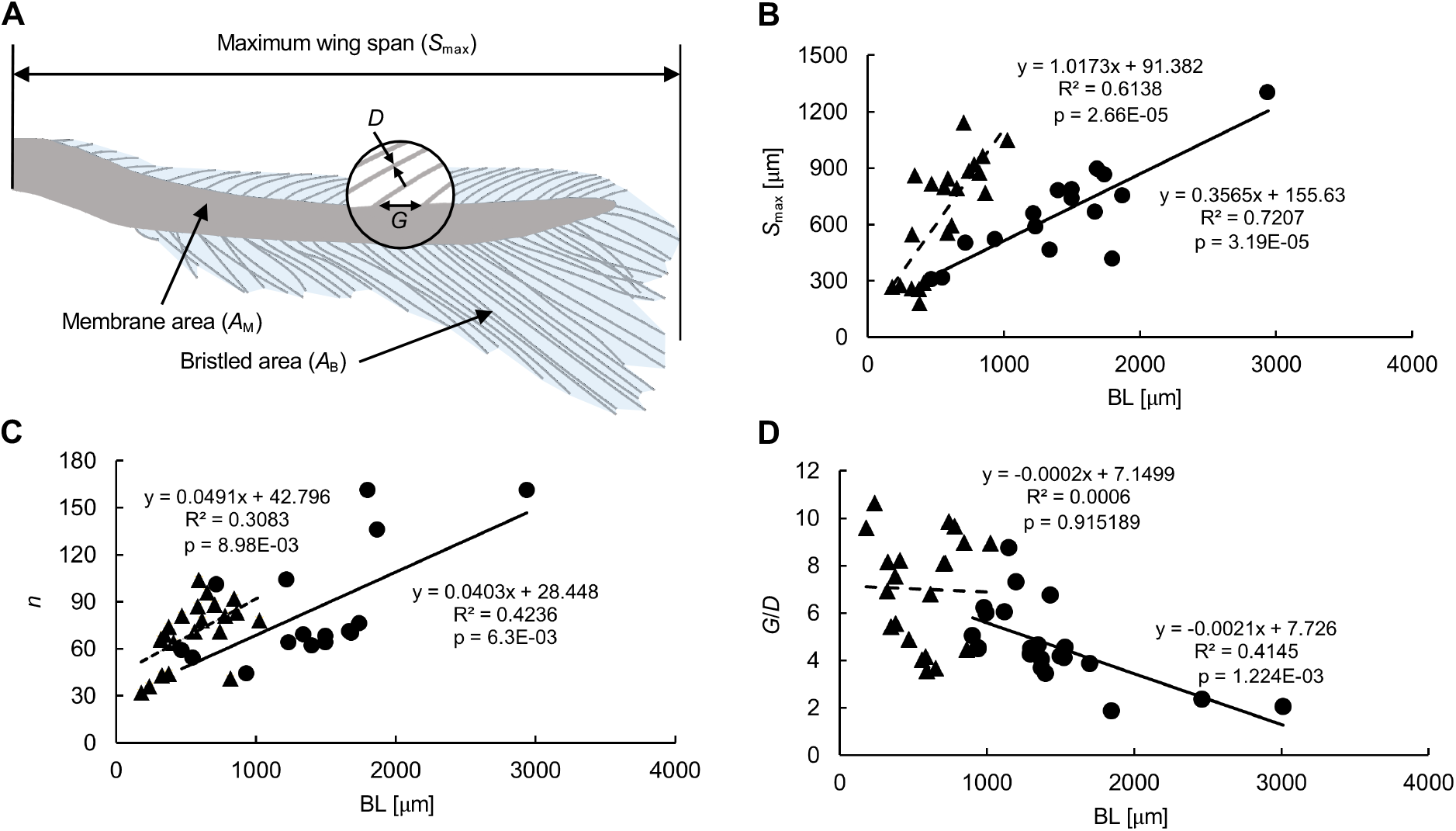
Morphological measurements of thrips (Thysanoptera) and fairyflies (Mymaridae) forewings. (A) Forewing of *Thrips setosus* (BL=1400 *μ*m) redrawn from Riley et al. (2011), with bristled area (*A*_B_), membrane area (*A*_M_), maximum wing span (*S*_max_), inter-bristle gap (*G*) and bristle diameter (*D*) indicated. (B) *S*_max_ as a function of BL (both in microns). (C) Number of bristles (*n*) as a function of BL. (D) *G/D* as a function of BL. Linear regressions for each data set are shown with *R^2^* and *p*-values. Fairyflies (---▲---); Thrips (--●--). The list of species used for *S*_max_ and *n* measurements are provided in Tables S1, S2. A different set of thrips forewing images were used for measuring *G/D* (see Table S3 for the list of species).

### Simplified wing models

Forewing morphological measurements in Thysanoptera and Mymaridae species showed a large variation of *n* (32 to 161). For a bristled wing of rectangular planform with constant *w* (Fig. 2A), *G* and *D*, *n* can be calculated using the following equation:

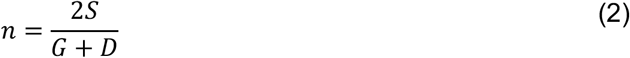

where *n* represents the total number of bristles on both sides of a solid membrane. We designed and fabricated 14 pairs of scaled-up, simplified (rectangular planform) physical wing models to examine effects of changing *G*, *D* and *S* (Table 1). In addition, 9 wing pairs were used to examine the variation in non-dimensional geometric variables: (i) *n* and (ii) *G/D* (Table 1). Note that we rounded down the *n* to a whole number in the physical models. As our wing models were scaled-up, we were not able to match *G*, *D* and *S* values to be in the range of tiny insects. To achieve geometric similarity, we maintained the relevant non-dimensional geometric variables (*n* and *G*/*D*) to be within their corresponding biological ranges in all the physical models.

**Figure 2.**
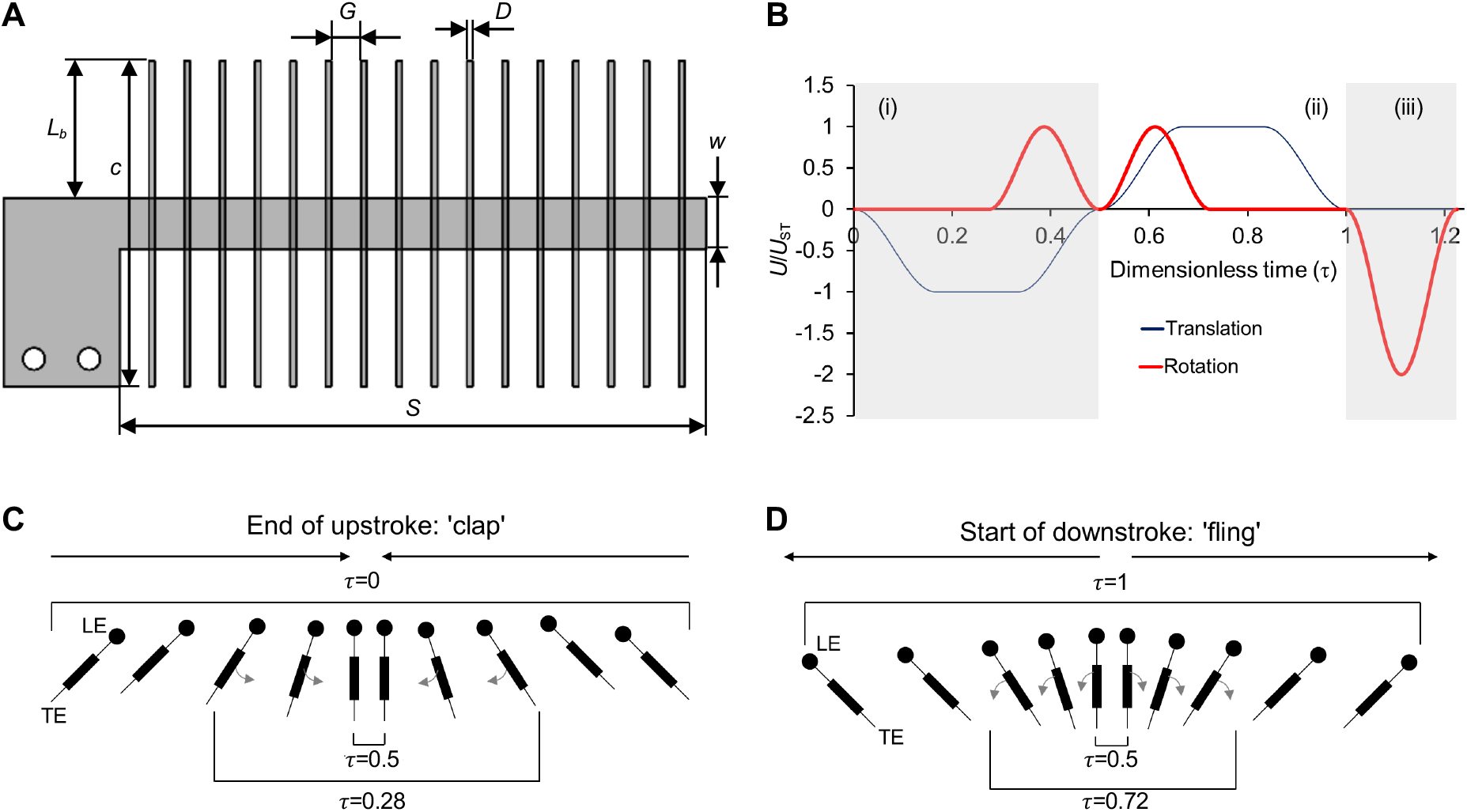
Physical bristled wing model and kinematics. (A) Diagram of the simplified bristled wing model with rectangular planform (*L*_b_=bristle length; *w*=membrane width). See Table 1 for the complete list of models tested. (B) Prescribed motion profile of a single wing, based on kinematics developed by Miller and Peskin (2005). Dimensionless velocity (*U*/*U*_ST_), is shown as a function of dimensionless time *τ* defined in Eqn 3. The wing motion consisted of rotation (thick line) and translation (thin line) along 3 regions: (i) clap (*τ*=0-0.5); (ii) fling (*τ*=0.5-1); (iii) 90-degrees wing rotation (*τ*=1-1.2) to position the wing for the start of the next cycle. During both clap and fling, wing translation was prescribed to occur throughout the wing rotation (100% overlap). The motion profiles prescribed to the other wing was identical in magnitude but opposite in sign, so that the wings would travel in opposite directions. Forces and PIV data were acquired from start of clap to the end of fling. Diagrammatic representation of wing motion during clap (C) and fling (D), where the sectional view along the wing span is shown. *τ* = 0, *τ* = 0.28, and *τ* = 0.5 correspond to start of clap (wings translating toward each other), start of wing rotation and end of clap, respectively. *τ* = 0.5, *τ* = 0.72, and *τ* = 1 correspond to start of fling with wings rotating and translating apart, end of wing rotation and end of fling, respectively. *U=*instantaneous wing tip velocity; *U*_ST_= steady translational velocity; LE=leading edge; TE=trailing edge.

**Table 1.**
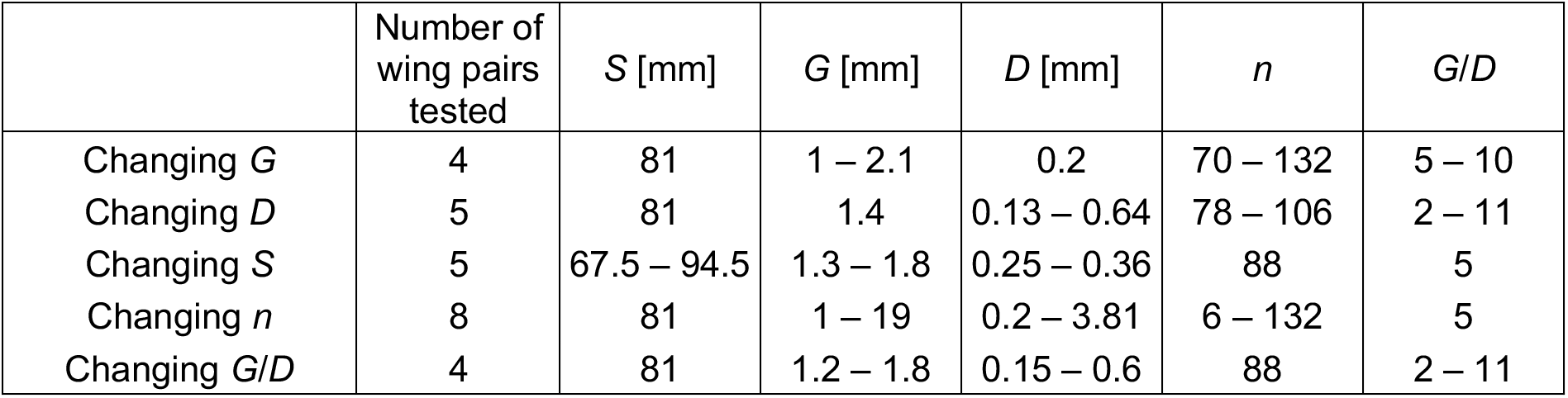
Experimental conditions and physical wing models used in this study. Each row represents the specific geometric variable or ratio that was controllably changed. Wing chord (*c*)=45 mm, membrane width (*w*)=7 mm, and bristle length (*L*_b_)=19 mm were maintained constant across all wing models. *G, D, S* and *n* represents inter-bristle gap, bristle diameter, wing span and number of bristles, respectively. 23 pairs of physical wing models were tested in this study. 3 wing pairs included in the case of varying *n* overlapped with 3 of the wing pairs considered in varying *D*, varying *S* and varying *G/D* conditions.

| | Number of<br>wing pairs<br>tested | $S$ [mm] | $G$ [mm] | $D$ [mm] | $n$ | $G/D$ |
| --- | --- | --- | --- | --- | --- | --- |
| Changing $G$ | 4 | 81 | 1 – 2.1 | 0.2 | 70 – 132 | 5 – 10 |
| Changing $D$ | 5 | 81 | 1.4 | 0.13 – 0.64 | 78 – 106 | 2 – 11 |
| Changing $S$ | 5 | 67.5 – 94.5 | 1.3 – 1.8 | 0.25 – 0.36 | 88 | 5 |
| Changing $n$ | 8 | 81 | 1 – 19 | 0.2 – 3.81 | 6 – 132 | 5 |
| Changing $G/D$ | 4 | 81 | 1.2 – 1.8 | 0.15 – 0.6 | 88 | 2 – 11 |

The bristled wings tested in this study were simplified to rectangular shape with constant wing chord (*c* in Fig. 2A) to minimize variability in confinement effects along the wing span from the tank walls. The percentage of *A*_M_/*A*_T_ in all the models was maintained at 15%, which is in the range of *A*_M_/*A*_T_ of thrips and fairyflies (Ford et al., 2019). Bristle length (*L*_b_, see Fig. 2A) on either side of the membrane as well as *w* were maintained as constants for all 23 wing models tested. The values of constants *c*, *L*_b_ and *w* are provided in Table 1.

The wing models were fitted into our robotic platform capable of mimicking the clap-and-fling kinematics. The 3 mm thick solid membrane used in all the wing models were 3D printed with polylactic acid (PLA) filament using Craftbot printers (CraftUnique LLC, Stillwater, OK, USA). The bristles were made of 304 stainless steel wires of varying diameter (Table 1), glued on top of the membrane. For flow visualization measurements using particle image velocimetry (PIV), we made new wing models with the solid membrane laser cut from 3 mm thick acrylic sheets. Also, to avoid reflection in PIV measurements, the bristles were blackened using a blackener kit (Insta-Blak SS-370, Electrochemical Products, Inc., New Berlin, WI, USA).

### Dynamically scaled robotic platform

The dynamically scaled robotic platform used in this study (Fig. 3A,B) has been described in previous studies (Kasoju et al., 2018, Ford et al., 2019) and experimentally validated against results in Sunada et al. (2002) corresponding to a single wing in translation at varying angles of attack (in Kasoju et al., 2018). Bristled wing models were attached to 6.35 mm diameter stainless steel D-shafts via custom aluminum L-brackets. Two 2-phase hybrid stepper motors with integrated encoders (ST234E, National Instruments Corporation, Austin, TX, USA) were used on each wing to perform rotation and translation. Rotational motion on a wing was achieved using a bevel gear for coupling a motor to a D-shaft. Translational motion was achieved using a rack and pinion mechanism driven by a second motor. All four stepper motors (for a wing pair) were controlled using a multi-axis controller (PCI-7350, National Instruments Corporation, Austin, TX, USA) via custom programs written in LabVIEW software (National Instruments Corporation, Austin, TX, USA). The assembly was mounted on an acrylic tank measuring 0.51 m x 0.51 m in cross-section, and 0.41 m in height. The tank was filled to 0.31 m in height with a 99% glycerin solution, such that the wings were completely immersed in the fluid medium.

**Figure 3.**
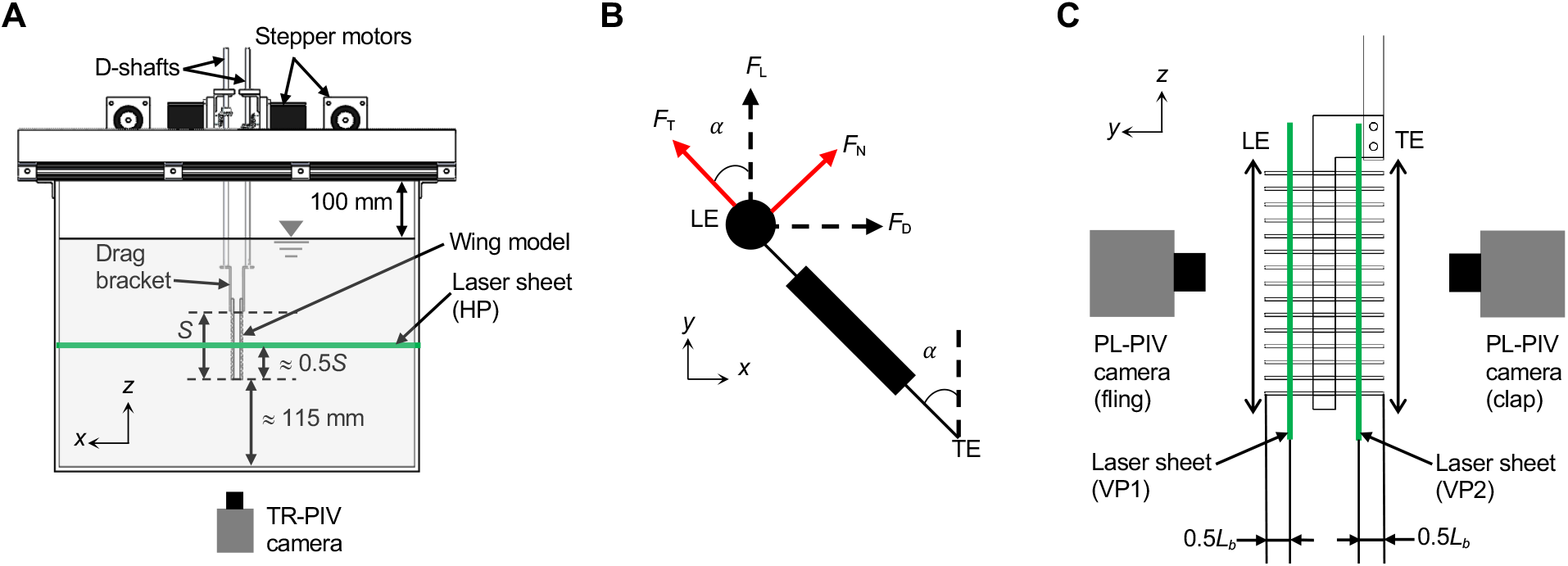
Robotic platform and experimental setup. (A) Front view of the robotic platform with bristled wings attached using custom L-brackets with strain gauges to measure the forces generated by a wing during clap and fling phases. The tank measured 510 mm x 510 mm in cross-section and 410 mm in height. 2D TR-PIV was used to visualize the chordwise flow field generated during clap and fling phases, where raw images were acquired using a high-speed camera and illumination was provided with a horizontally oriented laser sheet (horizontal plane, labeled HP) located approximately at mid-span (0.5*S*). (B) Sectional view along spanwise direction for a single bristled wing with directions of measured tangential (*F*_T_) and normal forces (*F*_N_) on a wing during rotation by angle *α* with respect to the vertical. Lift (*F*_L_) and drag (*F*_D_) forces were measured using a lift and drag bracket, respectively, by taking components of *F*_T_ and *F*_N_ in the vertical (*F*_L_) and horizontal (*F*_D_) directions. (C) 2D PL-PIV was used to measure the inter-bristle flow for 6 equally spaced time points during clap (*τ*~0.13 to *τ*~0.44) using a vertically oriented laser sheet (vertical plane 1, labeled VP1) and 7 equally spaced time points during fling (*τ*~ 0.63 to *τ*~0.94) at laser sheet labeled VP2. Both VP1 and VP2 were located at 0.5*L*_b_ from the LE and TE, respectively. *x,y,z* are fixed coordinate definitions.

### Kinematics

Due to the lack of adequately resolved free-flight recordings for characterizing instantaneous wing kinematics of tiny insects, we used a modified version of 2D clap-and-fling kinematics developed by Miller and Peskin (2005). Similar or modified forms of these kinematics have been used in several other studies (Miller and Peskin, 2009; Santhanakrishnan et al., 2014; Arora et al., 2014; Jones et al., 2016; Kasoju et al., 2018; Ford et al., 2019). The simplified kinematics used here do not capture: (a) 3D flapping translation during downstroke and upstroke, and (b) wing rotation at the end of downstroke (‘supination’). Fig. 2B shows the motion profiles prescribed for a single wing, where dimensionless velocity (instantaneous wing tip velocity *U* divided by steady translational velocity *U*_ST_) is provided as a function of dimensionless time (τ) during rotational and translational motion. Dimensionless time (τ) was defined as:

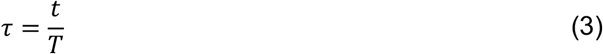

where *t* represents instantaneous time and *T* represents time taken to complete one cycle of clap-and-fling. The motion profile for the other wing was identical in magnitude but opposite in sign, so that the wings would travel in opposite directions. Both wings moved along a straight line (no change in elevation and stroke angles). Schematic diagrams of clap phase (Fig. 2C) and fling phase (Fig. 2D) are provided to show the direction of motion and wing position at the start and end of each portion of each half-stroke. The wings were programmed to start from an initial position corresponding to the start of the clap phase, and this was followed by the wings moving toward each other until the start of the fling phase after which the wings moved apart from each other. The distance between the wings at the end of clap phase was set to 10% of chord. The latter wing separation is similar to those observed in high-speed video recordings of freely flying thrips (Santhanakrishnan et al., 2014) and is also close enough to experience wing-wing interactions, but just far enough apart to prevent the leading and trailing edges of the rigid wing models from colliding during rotation. There was 100% overlap prescribed between rotation and translation during both clap and fling, meaning that the wings translated during the entire rotational time.

### Test conditions

Each wing model used in this study was tested at a chord-based Reynolds number of 10 (*Re_c_*=10). The kinematic viscosity (*v = μ/ρ*) of the 99% glycerin solution in which wing models were tested was measured using a Cannon-Fenske routine viscometer (size 400, Cannon Instrument Company, State College, PA, USA) to be 860 mm^2^ s^-1^ at standard room temperature. The chord-based Reynolds number was defined using the equation:

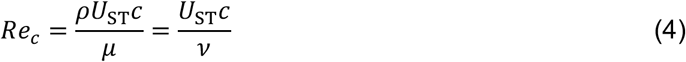

Using *Re_c_*=10 and the measured *v, U*_ST_ was calculated. Time-varying rotational and translational velocities were generated from the solved *U*_ST_ value using the equations in Miller and Peskin (2005). The complete duration of a clap and fling cycle (*T* in Eqn 3) was 2,220 ms. As *c* was not changed across all wing models (Table 1), *Re_c_* was constant for all wing models tested using the same motion profile.

### Force measurements

Similar to Kasoju et al. (2018) and Ford et al. (2019), force measurements were performed using L-brackets with strain gauges mounted in half-bridge configuration (drag bracket shown in Fig. 3A). The strain gauge conditioner continuously measured the force in form of voltage, and a data acquisition board (NI USB-6210, National Instruments Corporation, Austin, TX, USA) synchronously acquired the raw voltage data and angular position of the wings once a custom LabVIEW (National Instruments Corporation, Austin, TX, USA) program triggered to start the recording at the start of a cycle. Force data and angular position of the wings were acquired for complete duration of clap-and-fling motion (*τ*=0 to 1) at a sample rate of 10 kHz. We used the same processing procedures as in Kasoju et al. (2018) as briefly summarized here. The voltage signal was recorded prior to the start of motion for a baseline offset. To establish a periodic steady state in the tank, the setup was run for 10 consecutive cycles prior to recording the force data for 30 continuous cycles. The next step was to filter the raw voltage data in MATLAB (The Mathworks Inc., Natick, MA, USA) using a third order low-pass Butterworth filter with a cutoff frequency of 24 Hz. The baseline offset was averaged in time and subtracted from the filtered voltage data. The lift and drag brackets were calibrated manually, and the calibration was applied to the filtered voltage data obtained from the previous step to calculate forces. The forces that were calculated represent tangential (*F*_T_) and normal (*F*_N_) forces (Fig. 3B). The lift (*F*_L_) and drag (*F*_D_) forces acting on a wings were calculated using Eqns 5,6 given below:

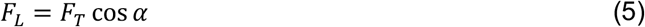

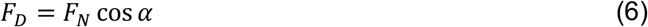

where *α* is the angular position of the wing relative to the vertical, recorded from the integrated encoder of the rotational stepper motor. Dimensionless lift coefficient (*C*_L_) and drag coefficient (*C*_D_) were calculated using the following relations:

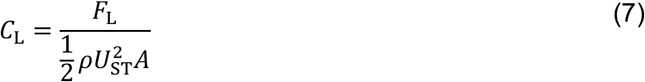

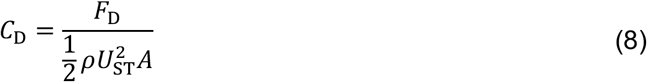

where *F*_L_ and *F*_D_ are the lift and drag forces (in Newtons), respectively, *ρ* is the fluid density (measured to be 1260 kg m^-3^), and *A* is the surface area of the rectangular planform of a wing (*A*=*S.c*). Standard deviations were calculated across 30 continuous cycles for *C*_L_ and *C*_D_, and the force coefficients were phase-averaged across all cycles to obtain time-variation of instantaneous force coefficients within a cycle. In addition, cycle-averaged force coefficients 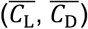 were calculated, with standard deviations and averages reported across 30 cycles for 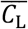 and 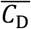. Note that all forces were only recorded on a single wing, with the assumption that forces generated by the other wing of a wing pair were equal in magnitude, as the motion was symmetric for both wings of a wing pair.

### Particle image velocimetry (PIV)

2D time-resolved PIV (2D TR-PIV) measurements were conducted to characterize the flow generated during clap-and-fling motion by bristle wing pairs along the chordwise plane (data acquired along a horizontal plane (HP) shown in Fig. 3A). 2D phase-locked PIV (2D PL-PIV) measurements were conducted to characterize flow leaked along the span of bristled wings (data acquired along 2 vertical planes (VP1 and VP2) shown in Fig. 3C).

#### 2D TR-PIV along wing chord

2D TR-PIV measurements were acquired for a total of 6 wing pairs, consisting of 2 wing pairs each for varying *G, D* and *S*. TR-PIV measurements were acquired along a chordwise (i.e. *x-y*) plane located at mid-span (Fig. 3A). The TR-PIV experimental setup and processing were similar to our previous studies (Kasoju et al., 2018; Ford et al., 2019) and is briefly summarized here. A single cavity Nd:YLF laser (Photonics Industries International, Inc., Bohemia, NY, USA) that provides a 0.5 mm diameter beam of 527 nm in wavelength was used in combination with a plano-concave cylindrical lens (focal length=-10 mm) to generate a thin laser sheet (thickness≈3-5 mm) positioned at mid-span (HP in Fig. 3A) to illuminate the field of view (FOV). TR-PIV images were acquired using a high-speed complementary metal-oxide-semiconductor (CMOS) camera with a spatial resolution of 1280 x 800 pixels, maximum frame rate of 1630 frames/s, and pixel size of 20 x 20 microns (Phantom Miro 110, Vision Research Inc., Wayne, NJ, USA). This camera was fitted with a 60 mm constant focal length lens (Nikon Micro Nikkor, Nikon Corporation, Tokyo, Japan). Hollow glass spheres of 10-micron diameter (110P8, LaVision GmbH, Göttingen, Germany) were used as seeding particles. A frame rate of 90 Hz was used to capture 100 evenly spaced images during both the clap and the fling phases. The raw images were processed using DaVis 8.3.0 software (LaVision GmbH, Göttingen, Germany) using one pass with an interrogation window of 64×64 pixels and two subsequent passes of 32×32 pixels window size. The processed TR-PIV images were phase-averaged over 5 cycles, and 2D velocity components and their positions were exported for calculating circulation (Γ) of the LEV and the trailing edge vortex (TEV). Γ was calculated for 8 equally spaced time points in both clap (from τ=0.05 to 0.4; increments of 5% of *τ*) and fling (from *τ*=0.55 to 0.9; increments of 5% of *τ*). Γ was calculated from the following equation using a custom MATLAB script:

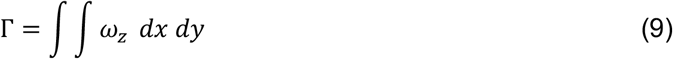

where *ω_z_* represents the out-of-plane (i.e., *z*) component of vorticity at leading or trailing edge, calculated from exported velocity vectors similar to Ford et al. (2019) and *dx dy* represents the area of the vorticity region selected for either the LEV or TEV. We used a minimum cutoff of 10% of the maximum of the overall maximum *ω_z_* at the leading and trailing edges for the time points tested. Γ was calculated for the right wing only, with the assumption that circulation for the left wing will be equivalent in magnitude but oppositely signed. Note that the left wing motion is symmetric to the right wing about *y-z* plane, making our assumption justifiable.

#### 2D PL-PIV along wingspan

The PL-PIV setup was similar to that used in Kasoju et al. (2018) and is briefly described here. Illumination was provided using a double-pulsed Nd:YAG laser (Gemini 200-15, New Wave Research, Fremont, CA) with a wavelength of 532 nm, maximum repetition rate of 15 Hz, and pulse width in the range of 3–5 ns. A 10 mm focal length cylindrical lens was used to generate a thin laser sheet (thickness≈3-5 mm) for illuminating the FOV. Raw PL-PIV images were acquired using a scientific CMOS (sCMOS) camera with a maximum spatial resolution of 2600 x 2200 pixels (maximum pixel size=6.5 x 6.5 microns) at a frame rate of 50 frames/s (LaVision Inc.,Ypsilanti, MI, USA), mounted with a 60 mm lens (same lens as in TR-PIV). The camera was focused on the seeding particles (same particles as in TR-PIV) along the laser sheet. PL-PIV measurements were acquired for all the wing models along 2 spanwise planes (VP1: fling and VP2: clap; see Fig. 3C) located at 0.5*L*b measured from the membrane. Raw image pairs were acquired at 6 time points in clap and 7 time points in fling, with adjacent time points spaced by 6.25% τ. Laser pulse separation intervals between the 2 images of an image pair ranged from 1,500 −19,831 *μ*s to obtain 6-8 pixels of particle displacement. The starting time point during clap phase (τ=0.0625) was neglected due to very small changes in flow surrounding the wings. For each wing model tested, 5 image pairs were acquired at each time point for 5 continuous cycles of clap and fling. The raw image pairs were processed using DaVis 8.3.0 using one pass with an interrogation window of 64 x 64 pixels and two subsequent passes of 32 x 32 pixels window size. The processed PL-PIV images were phase-averaged over 5 cycles and the velocity field was exported to quantify the amount of fluid leaked through the bristles along the wing span.

Cheer and Koehl (1987) proposed the use of a non-dimensional quantity called leakiness (*Le*) to characterize the amount of fluid leaking through bristled appendages. *Le* is defined as the ratio of the volumetric flow rate of fluid (*Q*) that is leaked through the inter-bristle gaps in the direction opposite to appendage motion under viscous conditions to that under inviscid conditions:

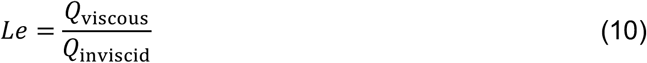

where *Q*_viscous_ represents the volumetric flow rate leaked through the bristles (i.e., opposite direction to wing motion) under viscous conditions, *Q*_inviscid_ represents the volumetric flow rate leaked through the bristles under no viscous forces (inviscid flow). Similar to Kasoju et al. (2018), we calculated the inviscid (or ideal) volumetric flow rate leaked through the bristles of a wing as:

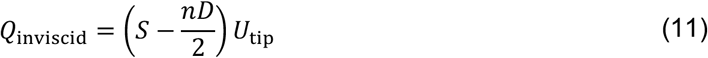

where *U*_tip_ represents wing tip velocity in the direction normal to the instantaneous wing position, defined as:

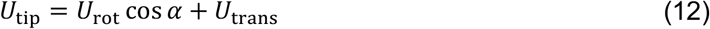

where *U*_trans_ and *U*_rot_ represent instantaneous translational and rotational velocities, respectively, and *α* represents instantaneous angle of a single wing relative to the vertical (Fig. 3B). *U*_rot_ was calculated as the product of the wing chord (*c*) and angular velocity of the wing (*ω*_rot_) as in Kasoju et al. (2018). *Q*_viscous_ was calculated from 2D PL-PIV velocity field data as the difference in volumetric flow rates of a solid (non-bristled) wing (denoted herein by *Q*_solid_) and the bristled wing under consideration, using the same steps as in Kasoju et al. (2018) that is also summarized here. 2D PL-PIV measurements were acquired on a solid wing model of the same *c* and *S* as that of the bristled wing under consideration, using identical motion profiles for both solid and bristled wings and at the same time points or ‘phase-locked’ positions. Horizontal velocity was extracted for the entire length of wingspan along a line ‘L’ that was oriented parallel to the wingspan and located downstream of the wing (i.e., in the direction of wing motion) at an *x*-distance of about 5% chord length from the rightmost edge of the wing surface when viewing the wing along the *x-z* plane. The horizontal component of the 2D PL-PIV velocity fields was in the direction normal to the wing, i.e., velocity component in the direction of wing motion. These velocity profiles were extracted for every wing model tested, at 6 time points in clap and 7 time points in fling. The viscous volumetric flow rate in the direction opposite to the wing motion (i.e., leaky flow) was calculated using the equation:

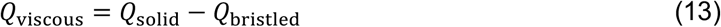

Volumetric flow rates (per unit width) for both solid and bristled wings about line ‘L’ was calculated by the line integral of the horizontal velocity using the equation below (in a custom MATLAB script):

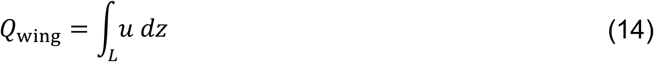

We did not directly estimate the reverse (i.e. leaky) viscous volumetric flow rate in the direction opposite to bristled wing motion from the 2D PL-PIV data due to the inability to simultaneously obtain high-magnification images needed to resolve flow through inter-bristle gaps (on the order of a few mm) along with lower magnification needed to resolve flow across the entire wing span (10x greater than *G*) for calculating *Q*_viscous_ across a bristled wing.

## RESULTS

### Forewing morphological analysis

For thrips and fairyflies, both *S*_max_ and *n* increased with increasing BL and showed strong positive correlation (Fig. 1B,C). For the 16 thrips species that were examined, *S*_max_ ranged from 305 *μ*m to 1301 *μ*m and *n* ranged from 44 to 161 (Table S1). For the 23 species of fairyflies that were examined, *S*_max_ ranged from 180 *μ*m to 1140 *μ*m and *n* ranged from 32 to 104 (Table S2). Values of *n* were found to be concentrated in the range of 30 to 90 for both thrips and fairyflies. Jones et al. (2016) reported that there was no correlation between the inter-bristle gap to bristle diameter ratio (*G*/*D*) and BL for fairyflies (Fig. 1D). By contrast, *G/D* for the 16 larger-sized thrips species examined were found to decrease with increasing BL and showed strong correlation (Fig. 1D, Table S3).

### Force measurements

For all the wing models tested, *C*_D_ and *C*_L_ were observed to follow the same trend in time during both clap and fling (Fig. 4A,B). Peak *C*_D_ occurred during fling (*τ*~0.6) in all wing models (Fig. 4A). This time point corresponds to end of rotational acceleration and translational acceleration (Fig. 2B), such that the wing pair would experience larger viscous resistance. *C*_D_ was found to drop after *τ*~0.6 until the wing rotation ended (*τ*~0.73) for all the wing models (Fig. 4A). Just before the *C*_D_ reached the negative value at the end of fling where the wings decelerate, we observed *C*_D_ to plateau from *τ*~0.73-0.84 (Fig. 4A). This time corresponds to steady translation motion of the wings (Fig. 2B), where the wings translate with constant velocity at 45° angle of attack (AOA). Most of the drag during a cycle was generated in fling. Time-variation of *C*_D_ was lower during clap half-stroke (*τ*=0-0.5) as compared to fling (Fig. 4A). Negative values of *C*_D_ during clap indicates that drag acts in the opposite direction as compared to drag force direction in fling.

**Figure 4.**
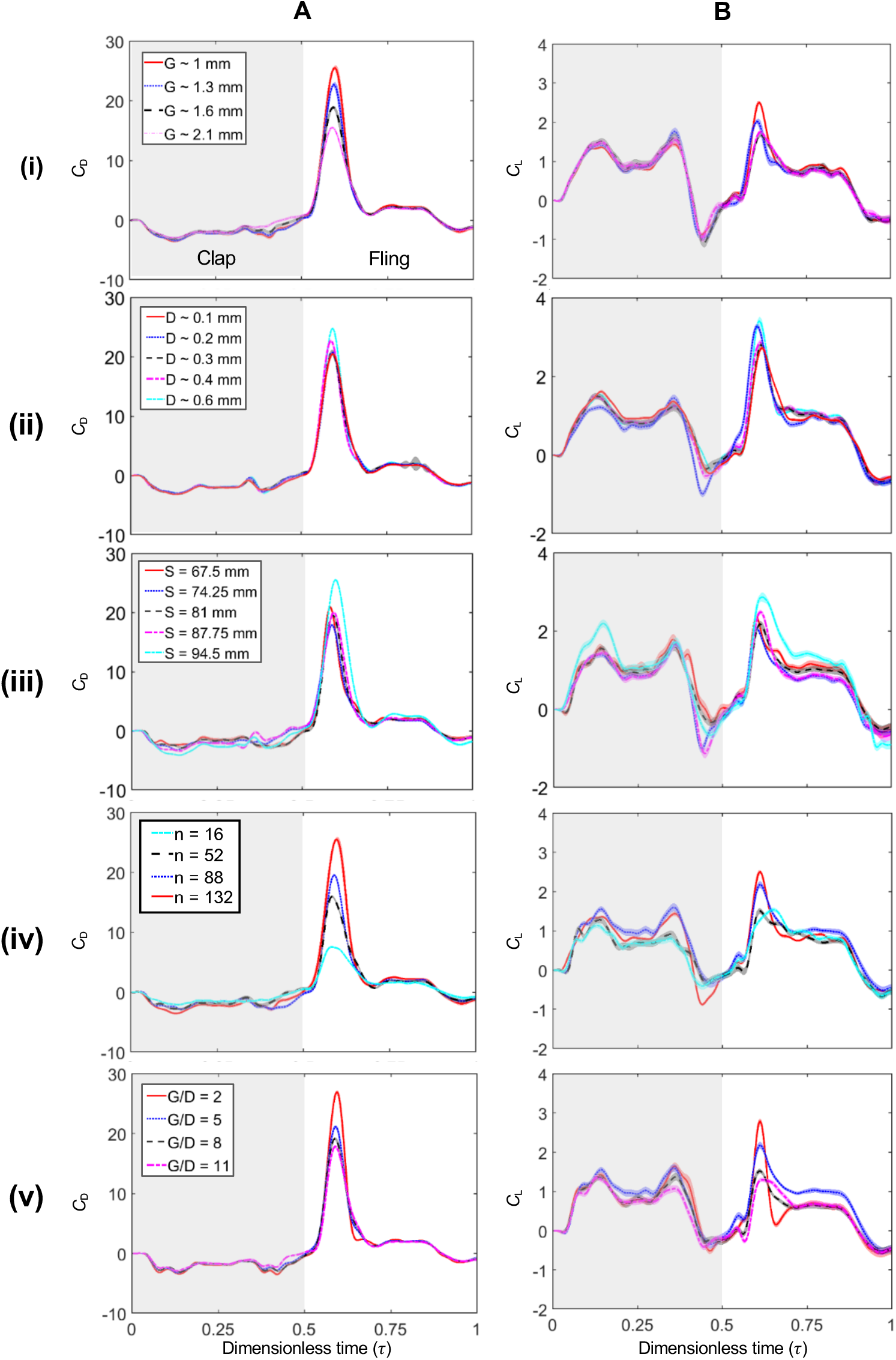
Time-varying force coefficients during clap and fling at *Re_c_*=10 with shading around each curve representing range of ±1 standard deviation (S.D) across 30 cycles. (A) and (B) show time-varying drag coefficient (*C*_D_) and lift coefficient (*C*_L_), respectively. From top to bottom, each row represents varying: (i) *G*, (ii) *D*, (iii) *S*, (iv) *n*, and (v) *G/D*. Gray shaded region in each plot represents the clap phase, while unshaded region represents the fling phase.

Three positive *C*_L_ spikes were observed in all the wing models (Fig. 4B): 1) τ~0.6 in fling, similar to that of peak *C*_D_; 2) start of clap (*τ*~0.16); and 3) end of clap (*τ*~0.38). *τ*~0.16 corresponds to the end of translational acceleration at 45° AOA and *τ*~0.38 corresponds to the end of rotational acceleration during clap (Fig. 2B). Peak *C*_L_ occurred during fling in majority of the wing models. Unlike the drag force, both clap and fling half-strokes contributed almost equally to lift generation.

Both *C*_D_ and *C*_L_ decreased with increasing *G* and decreasing *D* (Fig. 4(i),(ii)). Increasing *S* did not show any particular trend for *C*_D_ and *C*_L_ (Fig. 4(iii)). However, if we look at the extreme wingspans (67.5 mm and 94.5 mm), both *C*_D_ and *C*_L_ increased with increasing *S*. When increasing *n* for constant *G*/*D*, both *C*_D_ and *C*_L_ were found to increase (Fig. 4(iv)). In contrast, increasing *G/D* for constant *n* decreased both *C*_D_ and *C*_L_ (Fig. 4(v)).

Cycle-averaged force coefficients 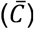 were used to examine how each geometric variable impacted aerodynamic forces in a complete cycle (Figs 5, 6). Individually increasing *G, D* and *S* showed negligible variation in 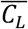 and 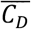 when considering the standard deviations (Fig. 5). 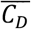 was found to increase with increasing *n* (Fig. 6A). Similarly, 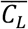 was found to increase until *n*=88 and then decreased with further increase in *n* (Fig. 6A). Interestingly, 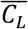 was found to be larger for *n* < 30 as compared to 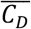, suggesting there may not be a particular, optimal *n* (i.e., largest 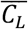 for smallest 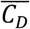) for *Re_c_*=10. Increasing *G/D* showed little to no variation in 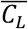 and 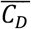 when considering the standard deviations (Fig. 6B).

**Figure 5.**
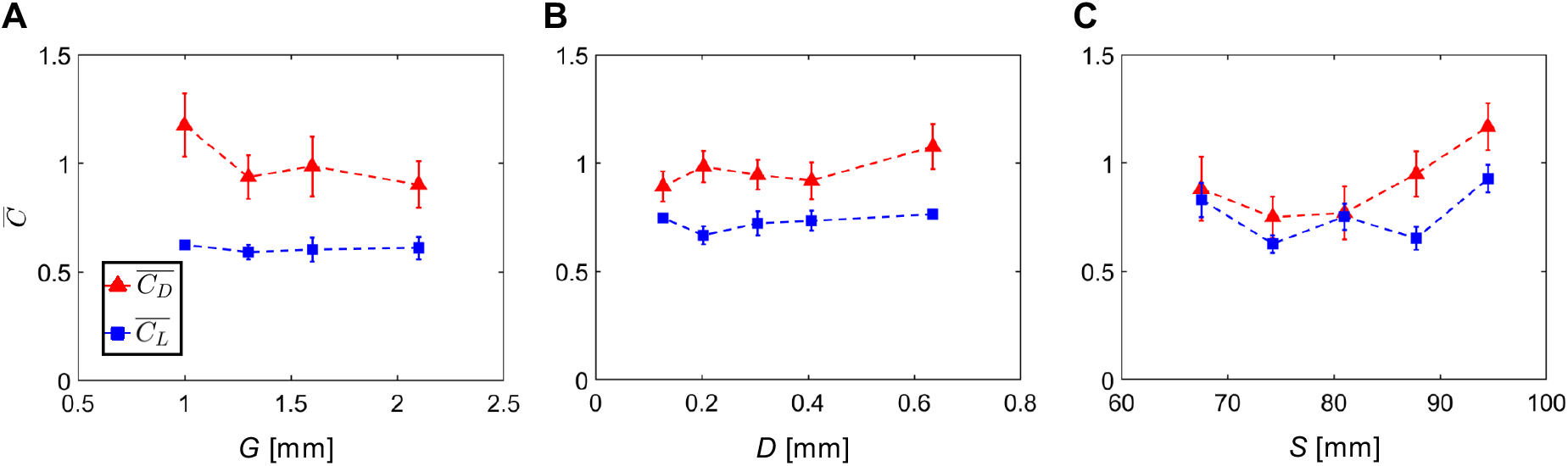
Cycle-averaged force coefficients 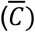 for varying *G, D* and *S*. Error bars corresponding to ±1 S.D are included for every datapoint. (A, B, C) show average lift coefficient 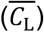 and average drag force coefficient 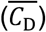 for varying *G, D*, and *S*, respectively. S.D estimates for 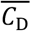 and 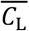 for all conditions were < 0.28 and < 0.1, respectively.

**Figure 6.**
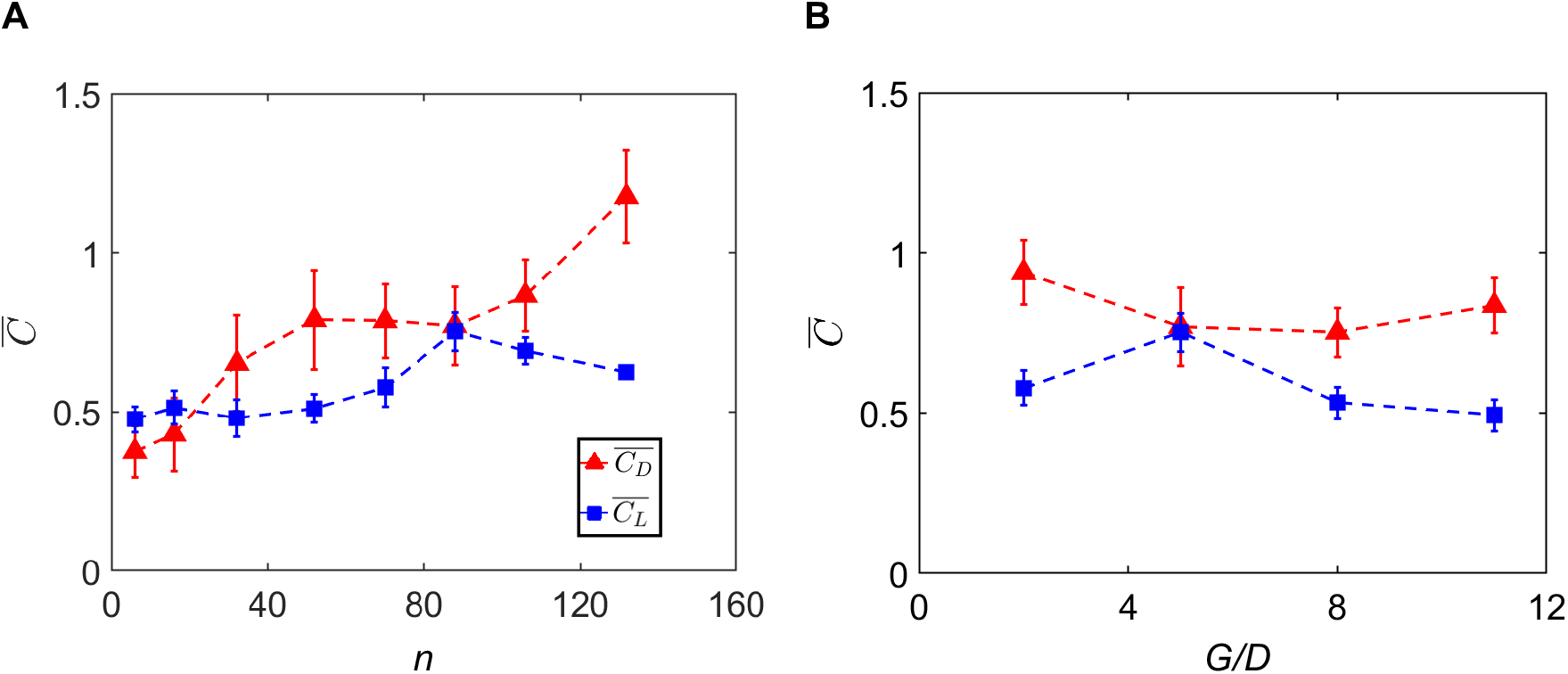
Cycle-averaged force coefficients 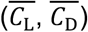 as a function of: (A) *n* and (B) *G*/*D*. Error bars corresponding to ±1 S.D are included. S.D estimates for 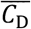 and 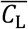 for all conditions were < 0.28 and < 0.1, respectively.

### Inter-bristle flow characteristics

Spanwise distribution of horizontal velocity (*u*) was examined near the time instant of peak *C*_D_ (*τ*~0.63) from 2D PL-PIV velocity fields (Fig. 7A). Looking at the extremes of each test condition, *u* increased with: (i) decreasing *G*; (ii) increasing *D*; (iii) increasing *S*; (iv) increasing *n*; and (v) decreasing *G*/*D*. This reveals how each variable (i.e., *G, D, S, n, G/D*) differentially affects flow through a bristled wing. *Le* was calculated using Eqn 10 and plotted in time (Fig. 7B). Similar to *C*_D_, *Le* was observed to peak during fling. During fling half-stroke, *Le* peaked either at *τ*~0.56 or *τ*~0.63 for all the wing models (Fig. 7B) where the wings were near the end of rotational acceleration (Fig. 2B). Similarly, wing deceleration during fling from *τ*~0.69 to *τ*~0.88 resulted in a drop in *Le* (Fig. 7B). During steady wing translation from *τ*~0.75 to *τ*~0.82, *Le* was found to almost plateau in all the wing models.

**Figure 7.**
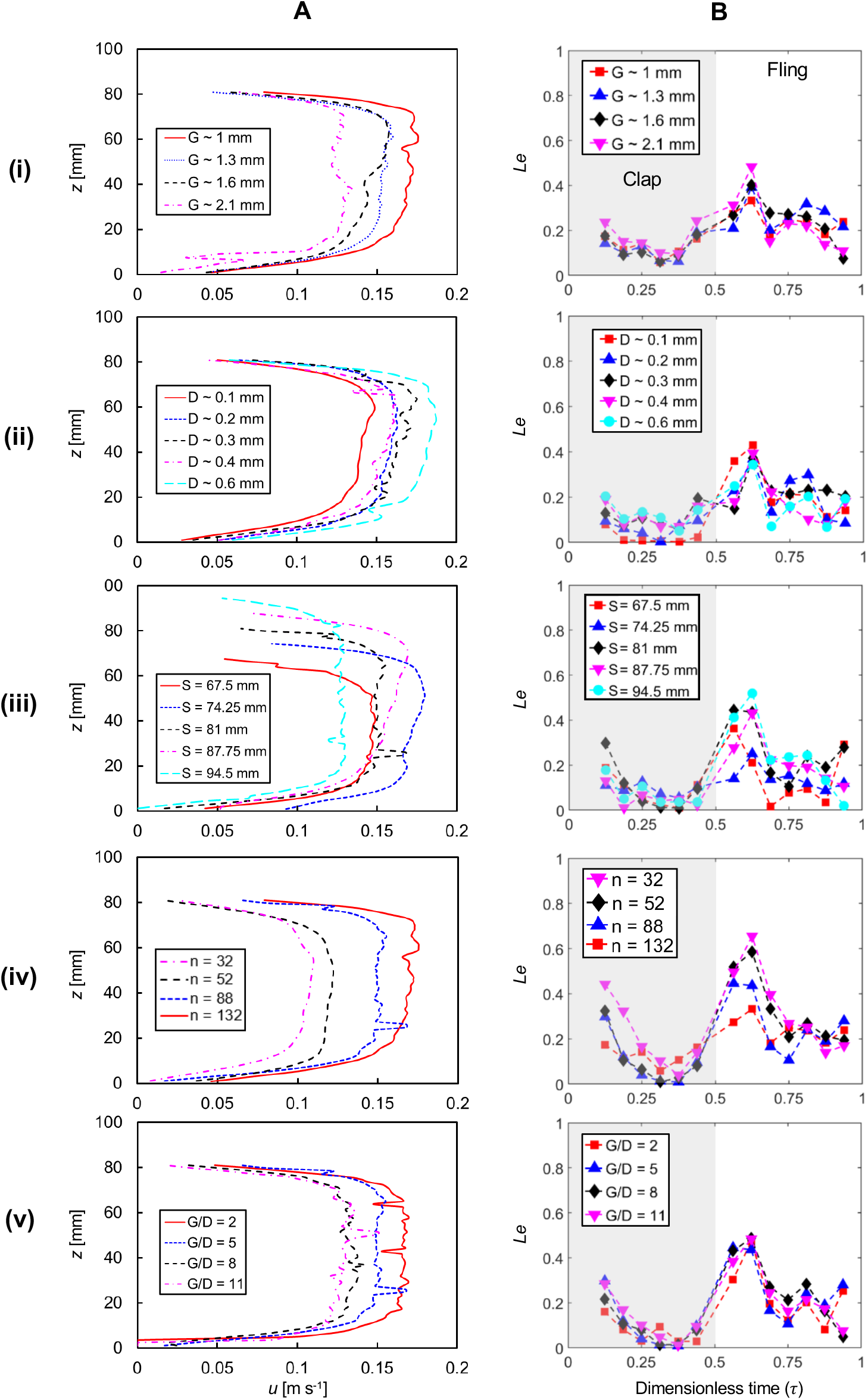
Inter-bristle flow characteristics. (A) Horizontal (i.e., *x*-component) velocity (*u*) variation along the wing span (*z*-direction) during fling at *τ*~0.63. The velocity profile was extracted at a vertical line *L* oriented parallel to the wing span, located at 5% chord length from the rightmost edge of the wing surface when viewing the wing along the *x-z* plane. (B) Time-variation of *Le*. From top to bottom, each row represents varying: (i) *G*, (ii) *D*, (iii) *S*, (iv) *n* and (v) *G/D*. Gray shaded region in column B represents the clap phase and unshaded region represents the fling phase.

*Le* was larger in early clap (*τ*~12.5) right after the wing pair just started from rest, with minimal time for boundary layers around each bristle to be well-developed. Thereafter, *Le* decreased with increasing clap duration until *τ*~0.38 corresponding to end of rotational acceleration (Fig. 2B). This latter observation in clap is in direct contrast to the peak in *Le* during fling that was observed at the end of rotational acceleration. This disparity can be explained by examining the prescribed wing motion. In clap, wings were prescribed to translate first at 45° AOA and then rotate. This provides ample time for the generation of shear layers around the bristles that block inter-bristle flow (see Kasoju et al., 2018 for a detailed discussion). Both rotation and translation started simultaneously in fling, necessitating more time for shear layers to develop around the bristles.

Peak *Le* increased with increasing *G* and decreasing *D* (Fig. 7B(i),(ii)). However, changes in *Le* were comparatively small for the range of variation in *G* and *D* tested in this study. Similar to force coefficients (Fig. 4(iii)), increasing *S* did not show any particular trend for *Le* (Fig. 7B(iii)). However, if we look at the extreme wingspans (67.5 mm and 94.5 mm), *Le* was found to increase with increasing *S*. Increasing *n* for constant *G/D* was found to decrease *Le*. Changing *G/D* for constant *n* showed little to no *Le* variation.

### Chordwise flow characteristics

Velocity vector fields overlaid on out-of-plane vorticity contours (*ω_z_*) showed the formation of LEV and TEV over the wing pair during clap and fling half-strokes (supplementary material Movies 1,2,3). Vorticity in the LEV and TEV increased near the end of clap and in early fling, when the wings were in close proximity of each other (Fig. 2C,D). This suggests that wing-wing interaction plays an important role in LEV and TEV formation, which in turn impacts force generation. Circulation (Γ) was calculated using Eqn 9 to quantify the strength of these flow structures. Γ of both the LEV and TEV showed little to no variation with changing *G, D* and *S*. Peak Γ for both the LEV and TEV occurred in fling (*τ*=0.65), near the end of both translational and rotational deceleration (Fig. 2B). This was followed by decrease in Γ of both LEV and TEV with increasing fling time (Fig. 8B,C,D). Γ of the LEV and TEV increased slowly in time during clap and reached a maximum near the end of the clap (*τ*=0.35), corresponding to the start of translational deceleration and end of rotational acceleration (latter being identical to the instant where peak Γ occurred in fling).

**Figure 8.**
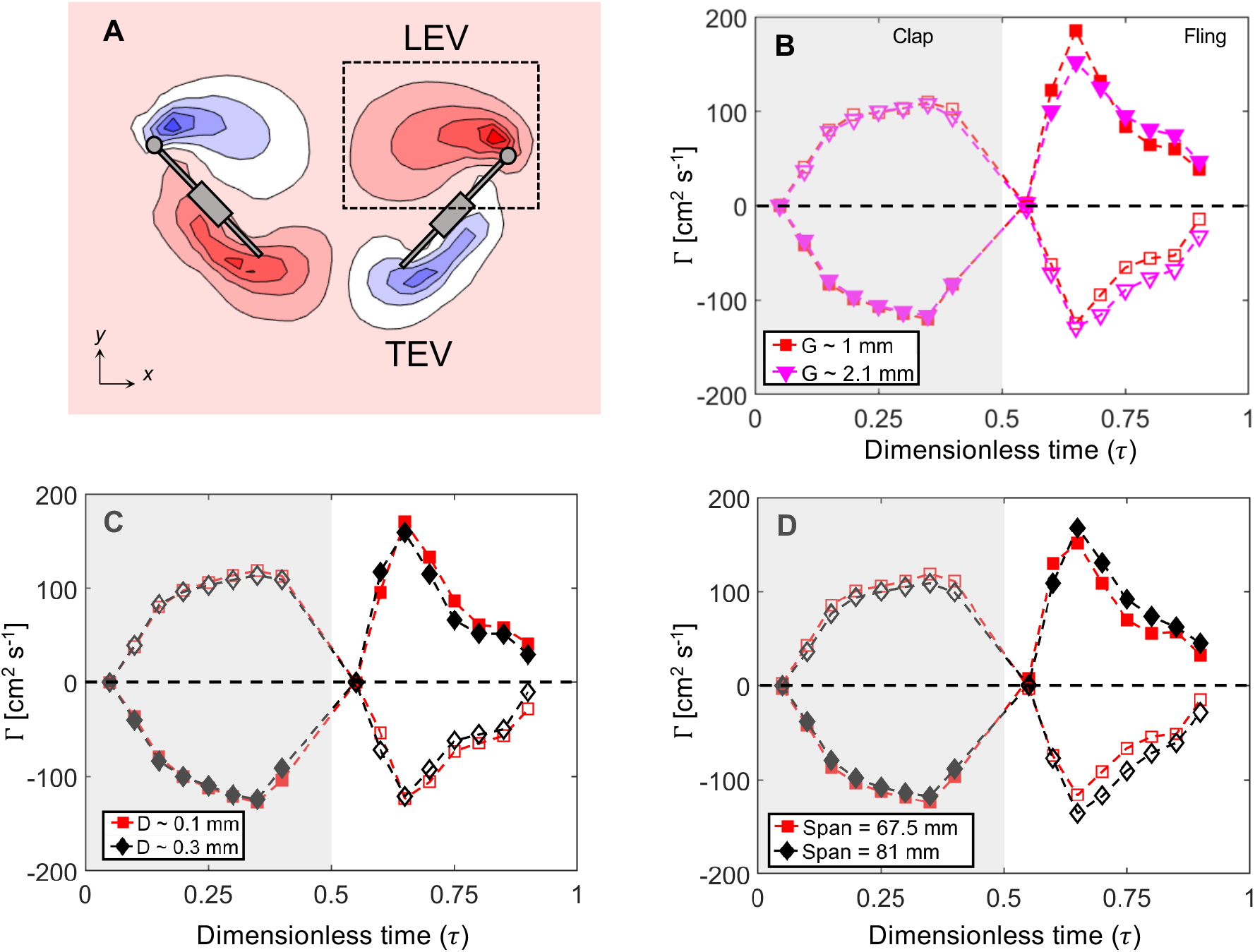
Chordwise flow and circulation (Γ). (A) Representative out-of-plane component of vorticity (*ω_z_*) during fling at *τ*=0.65, obtained from processed TR-PIV data. Γ about the right wing was calculated by drawing a box around the LEV and TEV separately and integrating *ω_z_* of the closed contour within each box. (B), (C) and (D) show Γ during clap and fling for varying *G, D* and *S*, respectively. Positive circulation corresponds to TEV during clap and LEV during fling. Negative circulation corresponds to LEV during clap and TEV during fling. Shaded markers represent circulation of LEV and hollow markers represents circulation of TEV.

From the prescribed kinematics (Fig. 2B), peak rotational acceleration starts early in fling, while it starts later into the clap. This could be the reason for Γ to peak early in fling and later in clap. This suggests that wing rotation plays a dominant role in LEV and TEV development. Also, both wings are in close proximity during the later stages of clap and early stages of fling, suggesting the importance of wing-wing interaction in in LEV and TEV development. Thus, wing rotation in concert with wing-wing interaction augments LEV and TEV circulation during both clap and fling half-strokes.

## DISCUSSION

Recent studies have shown that bristled wings provide drag reduction in clap-and-fling at *Re_c_* relevant to tiny insect flight (Santhanakrishnan et al., 2014; Jones et al., 2016; Kasoju et al., 2018; Ford et al., 2019). However, *n, S*_max_ and *G/D* have not been measured in different families of tiny insects, and their individual effects on aerodynamic forces are unclear. From analysis of forewings of 16 Thysanoptera (thrips) species consisting of 3 separate families and 21 Mymaridae (fairyflies) species, we found that *S*max and *n* were positively correlated with BL in both thrips and fairyflies. We also found that *G/D* in 22 species of thrips was negatively correlated with BL, in contrast to the lack of correlation between *G/D* and BL in fairyflies (Jones et al., 2016). Within the biologically relevant range of *n* and *G*/*D*, we find that: (1) increasing *G* provides more drag reduction as compared to decreasing *D*, (2) changing *n* for constant *G/D* has negligible impact on lift generation, and (3) changing *G/D* for constant *n* minimally impacts aerodynamic forces. The minimal influence of *n* on clap-and-fling aerodynamics, despite broad biological variation in *n* (32-161), suggests that tiny insects may experience lower biological pressure to functionally optimize *n* for a given wing span.

### Bristled wing morphology

Ford et al. (2019) reported a narrow range of *A*_M_/*A*_T_ (14%-27%) when examining the forewings of 25 thrips species. At *Re_c_* relevant to tiny insect flight, aerodynamic efficiency (lift-to-drag ratio) was found to be higher for *A*_M_/*A*_T_ in the range of thrips forewings. In this study, we measured *S*_max_, *n* and *G/D* in several species of Thysanoptera and Mymaridae. We found that both *S*_max_ and *n* on a wing increased with increasing BL in thrips and fairyflies (Fig. 1B,C). Interestingly, there was overlap in *S*_max_ (180-1301 *μ*m) across fairyflies and thrips. However, the majority of thrips species had BL > 1 mm as opposed to BL < 1 mm for all 21 fairyfly species. This suggests that there could be a limit to increasing wingspan in terms of aerodynamic performance. The values of *n* were concentrated in the range of 30-90 for the species of thrips and fairyflies that we examined. These observations led us to hypothesize that *n* may not need to be optimized to fall within a narrow range for a given wing span toward improving aerodynamic performance.

We also found that *G/D* negatively correlated with increasing BL in 16 species of thrips (Fig. 1D) unlike the lack of *G/D* to BL correlation in fairyflies reported by Jones et al., (2016). Previous studies (Jones et al., 2016; Kasoju et al., 2018) have reported that aerodynamic forces decrease with increasing *G*/*D*. The contrasting trend of *G/D* relative to BL between fairyflies and thrips raises a question as to whether *G/D* needs to be optimized across species for improving aerodynamic performance. However, it must be noted that we currently lack free-flight observations of fairyflies and thus do not know the extent to which they use flapping flight.

### Modeling considerations

Scaled-up physical models were used in this study to examine the roles of bristled wing geometric variables on clap-and-fling aerodynamics at *Re_c_*=10. We used this approach to overcome the difficulty of resolving the flow around and through a bristled wing of ~1 mm length. As we did not match the values of dimensional geometric variables to those of real insects, we used geometric similarity to match non-dimensional variables (*n*, *G*/*D*) in all the physical models to be in the range of tiny insects. As *n* depends on *G, D* and *S* per Eqn 2, the choices of non-dimensional variables include *n, G/D, G/S* and *D*/*S*. We chose to match *G/D* similar to Jones et al. (2016). In addition, to understand the isolated role of each dimensional variable, we tested scaled-up models varying in *G*, *D* and *S*. For each condition, we maintained the 2 other dimensional variables as constants and also matched the non-dimensional variables (*n, G/D*) to be within their biologically relevant ranges identified from morphological analysis.

Physical model studies of flapping flight match *Re_c_* of the experiments to biological values to achieve dynamic similarity. Specific to the bristled wings of interest to this study, dynamic similarity of inter-bristle flow characteristics also necessitates matching *Re_b_* to be in the range of tiny flying insects. When both *Re_c_* and *Re_b_* are matched between a physical bristled wing model to those of tiny insects, the scale model will produce similar non-dimensional forces to that of real insects. This is the major reason for presenting forces in term of non-dimensional coefficients throughout this study.

It has been reported that thrips (Kuethe, 1975) and *Encarsia Formosa* (Ellington, 1975) operate at *Re_b_*= 10^-2^ and 10^-1^, respectively and at *Re_c_* ~10. With the exception of Jones et al. (2016), the majority of modeling studies of bristled wing aerodynamics (Sunada et al., 2002; Santhanakrishnan et al., 2014; Lee and Kim 2017; Lee et al., 2018; Kasoju et al., 2018; Ford et al., 2019) only matched *Re_c_*~10 without matching *Re_b_* to be relevant to tiny insects. Matching *Re_b_* ensures that the flow through bristles of a model (and hence *Le*) would be similar to those of real insects. Considering that lift and drag are known to be impacted by the extent of leaky flow (Kasoju et al., 2018), we matched *Re_b_* to fall within 0.01 to 0.1 in majority of our physical models.

### Varying *G* and *D* for fixed *S*

Peak drag (*C*_D,max_) and lift (*C*_L,max_) coefficients were observed to generally increase with decreasing *G* and increasing *D*. However, changes in *C*_L,max_ when varying *G* or *D* (for fixed *S*) were substantially lower as compared to changes in *C*_D,max_, which was in agreement with our previous study on bristled wings with varying inter-bristle gap (Kasoju et al., 2018). Previous studies have proposed that substantial drag reduction realized with bristled wings in clap-and-fling is due to fluid leaking through the bristles (Santhanakrishnan et al., 2014; Jones et al., 2016; Kasoju et al., 2018). *Le* peaked at *τ*~0.56 or *τ*~0.63 (Fig. 7B) for each condition of varying *G* and varying *D*. Interestingly, both *C*_D,max_ and *C*_L,max_ were observed in between the same 2 time points, showing the importance of *Le* on dimensionless forces.

Previous studies of flow through bristled appendages have found that *Le* is a function of both *G* and *D* (Cheer and Koehl, 1987; Hansen and Tiselius, 1992; Leonard, 1992; Loudon et al., 1994). These studies also found that *Le* can be greatly influenced for *Re_b_* between 0.01 to 0.1, which is in the range of *Re_b_* for tiny insects. We calculated *Re_b_* for each wing model using *D* as the length scale in Eqn 4. *Re_b_* increases with increasing *D* and vice-versa. Within the biological *Re_b_* range (0.01-0.1), average force coefficients 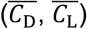 showed no variation when varying *D* (Fig. 9A,B). For varying *G*, we maintained *D* and *S* as constants. The calculated *Re_b_* for varying *G* tests was identical and within the biological *Re_b_* range. 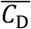 decreased with increasing *G* while 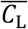 showed no variation (Fig. 9A,B).

**Figure 9.**
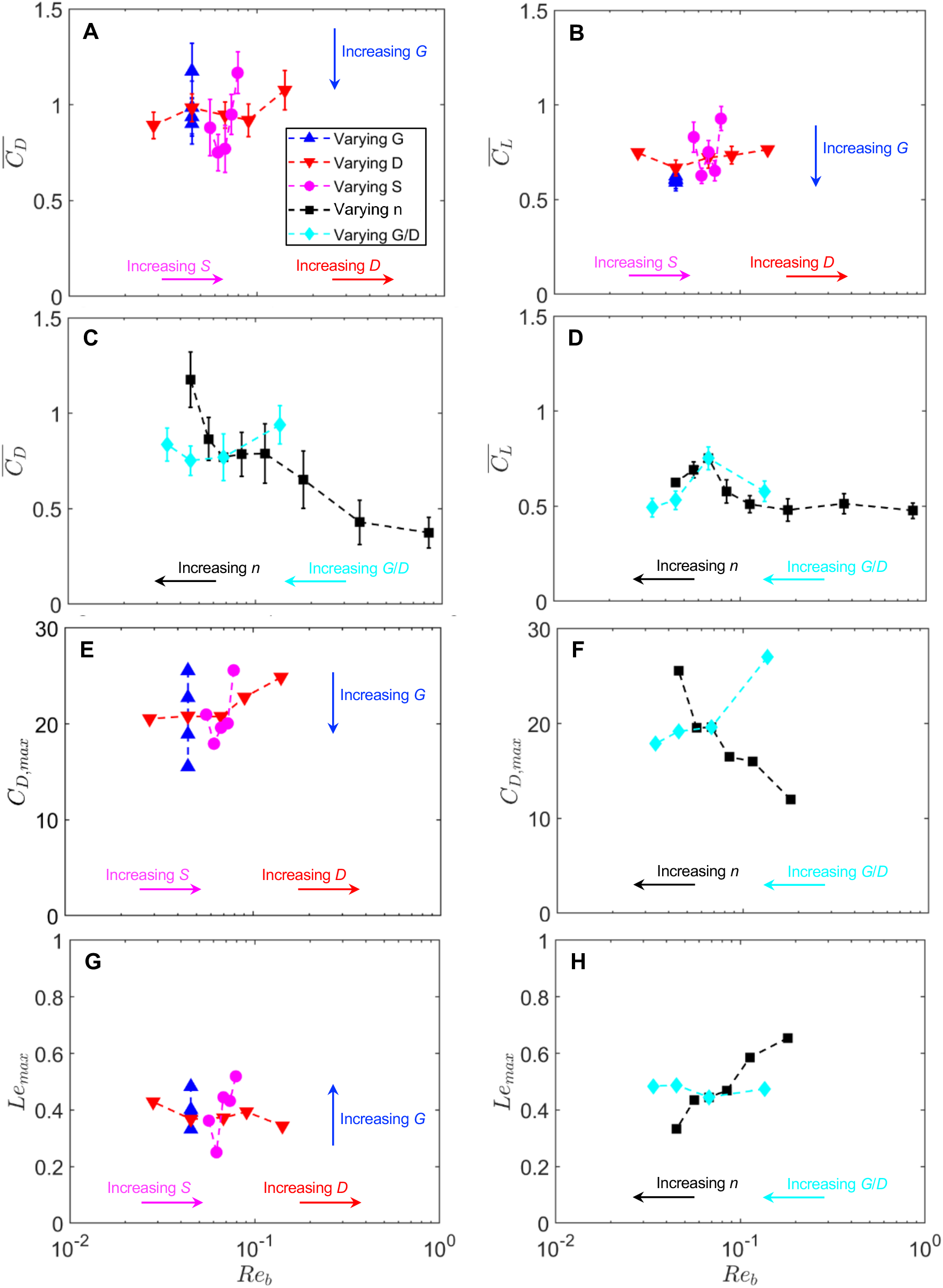
Average force coefficients 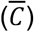, peak drag coefficient (*C*_D,max_) and peak leakiness (*Le*_max_) as a function of *Re_b_*. (A) and (B) show 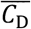 and 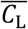, respectively, for varying *G, D* and *S*. (C) and (D) show 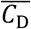 and 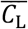, respectively, for varying *n* and varying *G*/*D*. (E) *C*_D,max_ for varying *G, D* and *S*. (F) *C*_D,max_ for varying *n* and *G*/*D*. (G) *Le*_max_ for varying *G, D* and *S*. (H) *Le*_max_ for varying *n* and *G/D*. *Re_b_* was calculated from Eqn 1 using bristle diameter (*D*) as the length scale. Trends with increasing geometric variables (*G, D, S, n*) and ratio (*G/D*) are indicated.

Increasing *Re_b_* via varying *D* showed opposite trends in *C*_D,max_ and *Le*_max_ (Fig. 9E,G). Within the biological *Re_b_* range, increasing *D* decreased *Le*_max_ and increased *C*_D,max_. Similarly, for a constant *Re_b_*, increasing *G* increased *Le*_max_ and decreased *C*_D,max_. These changes in leakiness for varying *G* and varying *D* are in agreement with previous studies (Cheer and Koehl,1987; Loudon et al., 1994). Further, for *Re_b_* in the range of 0.01-0.1, varying *G* showed larger changes in peak drag coefficients compared to varying *D*. Collectively, for *Re_b_* in the range of tiny insects, we find that varying *G* provides drag reduction (*C*_D,max_ and 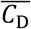) as compared to varying *D*, by augmenting *Le*. Tiny insects could possibly meet their flight demands by modulating the inter-bristle gap. Ellington (1980) observed that the dandelion thrips (*Thrips physapus*) open their forewing setae prior to takeoff, suggesting modulation of *G* may be possible when preparing for flight.

Little to no variation in 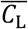 for both conditions (varying *G* and varying *D*) is attributed to formation of shear layers around the bristles that lowers the effective gap, resulting in the bristled wing behaving like a solid wing (Lee and Kim, 2017; Kasoju et al., 2018). Miller and Peskin (2005) proposed that LEV-TEV asymmetry plays a critical role in lift generation in clap-and-fling at *Re_c_* ~10. For varying *G* and varying *D*, we observed LEV circulation (Γ_LEV_) to be larger compared to TEV circulation (Γ_TEV_) for most of the clap-and-fling cycle (Fig. 8B,C). The implication of this asymmetry on lift generation can be seen by examining time-variation of *C*_L_ (Fig. 4B(i),B(ii)), where positive *C*_L_ was observed for most of the cycle. Both Γ_LEV_ and Γ_TEV_ peaked at *τ*=0.65, which corresponds to the same time point where peak *C*_L_ was observed. Minimal changes were observed in LEV-TEV asymmetry when increasing *G* and increasing *D* (compare Γ_LEV_-Γ_TEV_ in each case) resulting in little to no changes in *C*_L_ (Fig. 4B(i),B(ii)).

### Varying *S* for fixed *n* and *G/D*

Several studies examining the aerodynamic effects of varying *S* have reported contradictory findings. While some studies found little variation in force coefficients (Usherwood & Ellington, 2002; Luo & Sun, 2005; Garmann & Visbal, 2014), others have postulated that larger wing spans are detrimental for force generation (Harbig et al., 2012; Han, Chang & Cho, 2015; Bhat et al., 2019). All these studies considered solid wings at *Re_c_* >100. Our study is the first to report the effect of varying *S* on the aerodynamic performance of bristled wings performing clap-and-fling at *Re_c_*=10. Within the biological *Re_b_* range, more changes in 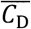 were observed when varying *S* as compared to 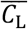 (Fig. 9A,B). In addition, *C*_D,max_ and *Le*_max_ increased with increasing *S* (Fig. 9E,G). Note that we varied *S* while maintaining *n* and *G/D* constant (*n*=88, *G*/*D*=5). To fit the same number of bristles while increasing *S*, we increased both *G* and *D* such that *G/D* was unchanged.

As previously discussed, changes in *D* within the biological *Re_b_* range produced negligible changes in force coefficients. The increase in *G* when increasing *S* is expected to increase *Le* and lower drag. However, we found that increasing *S* increased both *Le* and drag. Increasing *S* increases the wing surface area, which can explain the increase in drag. In addition, increasing *G* also increases *Le*. We speculate that the increase in *Le* with increasing *S* would minimize the increase in drag that would be expected from increasing wing surface area. Separately, varying *S* showed little changes in Γ_LEV_ and Γ_TEV_ (Fig. 8D) which resulted in small changes in *C*_L_ (Fig. 4B(iii)). Within the biological range of *n, G/D* and *Re_b_*, we postulate that larger *S* can be particularly beneficial to tiny insects when parachuting (Santhanakrishnan et al., 2014) as larger drag can slow their descent.

### Varying *n* for fixed *G/D* and S

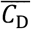 substantially increased with increasing *n* for a constant *G*/*D*, while 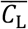 increased with *n* until *n*=88 and then decreased slightly with further increase in *n* (Fig. 6A). Wing models with *n*≤88 showed better aerodynamic performance in terms of force generation as compared to *n*>88. Interestingly, forewing morphological analysis showed that values of *n* were concentrated in the region 30-90 for thrips and fairyflies. 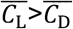 for bristled wing models with *n*=6 and 16, which can be interpreted as improved aerodynamic efficiency in flapping flight. Thrips have been observed to intermittently parachute (Santhanakrishnan et al., 2014), likely to lower the energetic demands of flapping flight and potentially also during wind-assisted long-distance dispersals. During parachuting, larger drag forces can assist them in migrating longer distances (Morse and Hoddle, 2006). 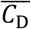 generated for *n* ranging between 30-90 (range of *n* for majority of the species considered here) was larger than 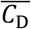 generated for *n*=6 and 16. In addition, our morphological measurements showed that *n* varied from 32-161, which can assist in generating lift needed for active flight as well as in generating drag needed for passive dispersal via parachuting.

Large variation in *C*_D,max_ and *Le*_max_ with *n* (Fig. 9F,H) shows the influence of the number of bristles on aerodynamic performance. *Le*_max_ was found to decrease with increasing *n*, while *C*_D,max_ was found to increase with increasing *n*. This suggests that changing *n* can aid or hinder aerodynamic performance by altering the leaky flow through the bristles. When varying *n* within the biological *Re_b_* range, only marginal changes in 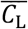 and 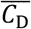 were observed (Fig. 9C,D). This suggests that for a fixed *S* and *G*/*D*, tiny insects may experience reduced biological pressure to fit a particular number of bristles for adequate lift generation. This inference is also supported by the broad inter-species variation in *n* (Fig. 1C).

### Varying *G*/*D* for fixed *n* and S

Little to no variation in 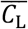 and 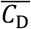 was observed when varying *G/D*, particularly when considering the standard deviations (Fig. 6B). Within the biological *Re_b_* range, *C*_D,max_ and *Le*_max_ were found to minimally change with increasing *G/D* (Fig. 9F,H). Also, varying *G/D* within the biological *Re_b_* range produced little to no variation in 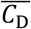 and 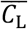. Note that for varying *G/D* within the biological *Re_b_* range, the inter-bristle gap in the corresponding physical models was nearly identical. From these results, we summarize that within the biological range of *Re_b_, G/D* variation for a fixed *S, n* and *G* result in little variation in aerodynamic force generation.

Interestingly, morphological measurements showed that *G/D* in thrips decreased with increasing BL, while no correlation between *G/D* and BL was reported for fairyflies (Jones et al., 2016). This dissimilar behavior in fairyflies and thrips raises a question regarding our use of static wing images for *G/D* measurements as opposed to free-flight wing images. We were restricted to using static forewing images due to the lack of free-flight wing images of tiny insects with adequate (i.e., high) magnification. It is unknown at present whether tiny insects can modulate *G/D* during free-flight, as such a capability can permit them to tailor aerodynamic forces in relation to ambient conditions (e.g., temperature, humidity, wind speed) and energetic costs.

### Limitations

As we used published forewing images for morphological analysis, we were unable to ascertain whether the positions of the bristles were unaffected during imaging. While we ensured that there was no visual damage to the bristles in the images that were used for analysis, it is possible that the measurements of *G* were affected by the above positioning uncertainty. We did not consider the effects of the following morphological variables: (a) asymmetry in *L*_b_ on either side of the forewing (i.e., leading edge and trailing edge); (b) angle of the bristles relative to the horizontal. It is possible that asymmetry in *L*_b_ within the biological *Re_b_* range does not noticeably impact clap-and-fling aerodynamics, as it is not unrealistic to expect damages to occur to the wing bristles during an insect’s life cycle. Similar to *G*, the angle of the bristles can be impacted during wing positioning for microscopy. High-magnification images of freely-flying tiny insect wings are needed to address these two measurement uncertainties. Finally, our physical models did not account for variation in wing shape and were simplified to a rectangular planform. There is tremendous diversity in wing shape, especially when comparing thrips (smaller chord relative to span) to fairyflies (teardrop-shaped). Future studies are needed to document inter-species diversity in wing shape and examine how they impact aerodynamic forces.

## Supporting information

Tables S1,S2,S3; Captions for Supplementary Movies 1 to 3

## LIST OF SYMBOLS AND ABBREVIATIONS

*α*: instantaneous angle of the wing relative to the vertical
Γ: circulation of a vortex
Γ_LEV_: circulation of the leading-edge vortex
Γ_TEV_: circulation of the trailing-edge vortex
*μ*: dynamic viscosity of fluid
*v*: kinematic viscosity of fluid
*ρ*: fluid density
*τ*: dimensionless time
*ω_z_*: *z*-component of vorticity
*A*: surface area of rectangular planform wing
*A*_B_: area occupied by bristles of a bristled wing
*A*_M_: area of solid membrane of a bristled wing
*A*_T_: total wing area
AOA: angle of attack
BL: body length
*c*: wing chord
*c*_ave_: average wing chord
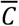: cycle-averaged force coefficient
*C*_D_: drag coefficient
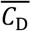: cycle-averaged drag coefficient
*C*_D,max_: peak drag coefficient
*C*_L_: lift coefficient
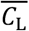: cycle-averaged lift coefficient
*C*_L,max_: peak lift coefficient
CMOS complementary: metal-oxide-semiconductor
*D*: bristle diameter
*F*_T_: tangential force on a wing
*F*_N_: normal force on a wing
*F*_D_: drag force
*F*_L_: lift force
FOV: field of view
*G*: inter-bristle spacing (or gap)
*G*/*D*: inter-bristle gap to bristle diameter ratio
HP: horizontal plane
*L*_b_: bristle length on either side of the solid membrane of a bristled wing
*Le*: leakiness
*Le*_max_: peak leakiness
LEV: leading edge vortex
*n*: number of bristles
PIV: particle image velocimetry
PLA: polylactic acid
PL-PIV: phase-locked PIV
*Q*: volumetric flow rate of fluid
*Q*_bristied_: *Q* for bristled wing
*Q*_inviscid_: volumetric flow rate leaked through the bristles under no viscous forces (inviscid flow)
*Q*_solid_: *Q* for solid wing
*Q*_viscous_: volumetric flow rate leaked through the bristles under viscous conditions
*Re*: Reynolds number
*Re_b_*: Reynolds number based on bristle diameter
*Re_c_*: Reynolds number based on wing chord
*S*: wing span of a rectangular wing
*S*_max_: maximum wing span
*t*: instantaneous time
*T*: time duration for one cycle of clap-and-fling
TEV: trailing edge vortex
TR-PIV: time-resolved PIV
*U*: instantaneous wing tip velocity
*U*_rot_: instantaneous rotational velocity
*U*_ST_: steady translational velocity
*U*_TIP_: wing tip velocity in the direction normal to the instantaneous wing position
*U*_trans_: instantaneous translational velocity
VP: vertical plane
*w*: membrane width

## ACKNOWLEDGEMENTS

None.

## COMPETING INTERESTS

The authors declare no competing or financial interests.

## AUTHOR CONTRIBUTIONS

Conceptualization: V.T.K. and A.S.; Methodology: V.T.K., M.P.F. and A.S.; Physical model fabrication: V.T.K. and T.T.N.; Image analysis, experimental data acquisition and processing: V.T.K., M.P.F. and T.T.N.; Data analysis and interpretation: V.T.K. and A.S.; Writing (original draft, review and editing): V.T.K., M.P.F. and A.S.; Project administration: A.S.; Funding acquisition: A.S.

## FUNDING

This work was supported by the National Science Foundation [CBET 1512071 to A.S.], Lew Wentz Foundation at Oklahoma State University [Wentz Research Grant to T.T.N.], and the College of Engineering, Architecture and Technology (CEAT) at Oklahoma State University [CEAT Undergraduate Research Scholarship to T.T.N.].

## DATA AVAILABILITY

Data generated from this study are included in the manuscript and supplementary material.

## Notes

### Competing Interest Statement

The authors have declared no competing interest.

