## Supplementary material for "Inter-species variation in number of bristles on forewings of tiny insects does not impact clap-and-fling aerodynamics": Tables S1,S2,S3; Captions for Supplementary Movies 1 to 3

*ELECTRONIC SUPPLEMENTARY MATERIAL*

### SUPPLEMENTARY TABLES

**Table S1.** Forewing morphological characteristics of thrips (Thysanoptera) species considered in this study. BL = body length; *n* = number of forewing bristles; $S_{max}$ = maximum wing span; $A_T$ = total wing area; $c_{ave}$ = average wing chord calculated as $A_T/S_{max}$.

| Species | BL [$\mu$m] | *n* | $S_{max}$ [$\mu$m] | $A_T$ [$\mu m^2$] | $c_{ave}$ [$\mu$m] | Source |
|---|---|---|---|---|---|---|
| *Gynaikothrips ficorum* | 1800 | 161 | 416 | 51271 | 123.25 | *MAF lab (2011) |
| *Hoplandothrips* sp. | 1870 | 136 | 753 | 179289 | 238.1 | *MAF lab (2011) |
| *Karnyothrips* sp. | 1340 | 69 | 462 | 72010 | 155.87 | *MAF lab (2011) |
| *Leucothrips piercei* | 550 | 54 | 315 | 31950 | 101.43 | *MAF lab (2011) |
| *Limothrips cerealium* | 1220 | 104 | 656 | 131038 | 199.76 | *MAF lab (2011) |
| *Neohydatothrips samayunkur* | 720 | 101 | 500 | 64836 | 129.68 | *MAF lab (2011) |
| *Scirtothrips* sp. | 470 | 59 | 305 | 26623 | 87.29 | *MAF lab (2011) |
| *Liothrips ludwigi* | 2940 | 161 | 1301 | 457976 | 352.02 | Zamar et al., 2013 |
| *Frankliniella occidentalis* | 1685 | 70 | 896 | 178471 | 199.19 | Riley et al., 2011 |
| *Thrips tabaci* | 1500 | 68 | 785 | 143076 | 182.27 | Riley et al., 2011 |
| *Frankliniella schultzei* | 1670 | 71 | 666 | 119750 | 179.81 | Riley et al., 2011 |
| *Scirtothrips dorsalis* | 935 | 44 | 518 | 71988 | 138.98 | Riley et al., 2011 |
| *Frankliniella intonsa* | 1740 | 76 | 863 | 141059 | 163.46 | Riley et al., 2011 |
| *Thrips setosus* | 1400 | 62 | 779 | 135609 | 174.09 | Riley et al., 2011 |
| *Ceratothripoides claratris* | 1235 | 64 | 587 | 84398 | 143.78 | Riley et al., 2011 |
| *Frankliniella gemina* | 1500 | 64 | 737 | 130100 | 176.53 | Riley et al., 2011 |

*MAF Plant Health & Environment Laboratory (2011)

**Table S2.** Forewing morphological characteristics of fairyfly (Mymaridae) species considered in this study. Definitions of symbols and abbreviations are in caption of Table S1.

| Species | BL [$\mu$m] | *n* | $S_{max}$ [$\mu$m] | $A_T$ [$\mu m^2$] | $c_{ave}$ [$\mu$m] | Source |
|---|---|---|---|---|---|---|
| *Eustochus nearticus* | 703.5 | 88 | 1140 | 301731 | 264.68 | Huber & Baquero, 2007 |
| *Eustochus besucheti* | 742 | 71 | 885 | 193588 | 218.75 | Huber & Baquero, 2007 |
| *Eustochus pengellyi* | 781 | 81 | 919 | 187288 | 203.8 | Huber & Baquero, 2007 |
| *Eustochus yoshimotoi* | 845 | 92 | 965 | 226967 | 235.2 | Huber & Baquero, 2007 |
| *Eustochus nipponicus* | 1023.5 | 78 | 1050 | 237216 | 225.92 | Huber & Baquero, 2007 |
| *Mymaromella pala* | 321.5 | 66 | 260 | 22306 | 85.8 | Huber et al., 2008 |
| *Mymaromella mira* | 376 | 74 | 256 | 23297 | 91.01 | Huber et al., 2008 |
| *Mymaromella chaoi* | 378 | 44 | 180 | 11406 | 63.37 | Huber et al., 2008 |
| *Mymaromella cyclopterus* | 409 | 64 | 290 | 27941 | 96.35 | Huber et al., 2008 |
| *Stethynium brevioviipositor* | 585 | 87 | 557 | 86512 | 155.32 | Huber et al., 2006 |
| *Stethynium ophelimi* | 590 | 104 | 843 | 178413 | 211.65 | Huber et al., 2006 |
| *Kikiki huna* | 180 | 32 | 266 | 20101 | 75.57 | Huber & Noyes, 2013 |
| *Tinkerbella nana* | 237.5 | 36 | 276 | 26113 | 94.62 | Huber & Noyes, 2013 |
| *Dicopomorpha schleideni* | 326.5 | 43 | 548 | 57316 | 104.6 | Lin et al., 2007 |
| *Mimalaptus victoria* | 346.5 | 69 | 862 | 237236 | 275.22 | Lin et al., 2007 |
| *Cleruchoides noackae* | 470.5 | 81 | 815 | 105608 | 129.59 | Lin et al., 2007 |
| *Prionaphes depressus* | 559.5 | 71 | 800 | 166574 | 208.22 | Lin et al., 2007 |
| *Schizophragma basalis* | 614 | 78 | 594 | 84870 | 142.88 | Lin et al., 2007 |
| *Allanagrus magniclava* | 653 | 96 | 793 | 137873 | 173.87 | Lin et al., 2007 |
| *Richteria lamennaisi* | 819 | 41 | 876 | 158410 | 180.83 | Lin et al., 2007 |
| *Eubroncus dubius* | 861 | 83 | 770 | 100113 | 130.02 | Lin et al., 2007 |

**Table S3.** Thrips (Thysanoptera) species considered in this study for measurement of inter-bristle gap (*G*) to bristle diameter (*D*) ratio. BL = body length.

| **Species** | **BL [$\mu$m]** | ***G*/*D*** | **Source** |
|---|---|---|---|
| *Yaobinthrips yangtzei* | 1500 | 4.19 | Zhang et al., 2010 |
| *Pandanothrips ryukyuensis* | 1530 | 4.57 | Masumoto & Okajima, 2013 |
| *Pandanothrips wangi* | 1120 | 6.05 | Masumoto & Okajima, 2013 |
| *Frankliniella veracrucensis* | 1295 | 4.27 | Goldaracena & Hance, 2017 |
| *Haplothrips herajius* | 1845 | 1.88 | Minaei & Aleosfoor, 2013 |
| *Akarethrips iotus* | 1525 | 4.11 | Dang et al., 2014 |
| *Frankliniella brunneicornis* | 2460 | 2.37 | Mound & Reynaud, 2005 |
| *Frankliniella strasseni* | 3010 | 2.05 | Mound & Reynaud, 2005 |
| *Pseudodendrothrips marissae* | 900 | 5.06 | Mound & Tree, 2016 |
| *Hydatothrips aliceae* | 1370 | 3.69 | Mound & Tree, 2009 |
| *Bhattithrips borealis* | 1370 | 4.05 | Mound, 2009 |
| *Siamothrips balteus* | 940 | 4.53 | Wang & Tang, 2016 |
| *Clypeothrips idrisi* | 1350 | 4.65 | Ng & Mound, 2015 |
| *Neohydatothrips clavisetis* | 980 | 6.23 | Lima & Mound, 2016 |
| *Neohydatothrips notialis* | 995 | 6.01 | Lima & Mound, 2016 |
| *Neohydatothrips renatae* | 940 | 4.51 | Lima & Mound, 2016 |
| *Lenkothrips kaminskii* | 1300 | 4.52 | Cavalleri & Mound, 2016 |
| *Haplothrips dissociates* | 1700 | 3.87 | Cavalleri et al., 2016 |
| *Kenyattathrips katarinae* | 1400 | 3.45 | Mound, 2009 |
| *Thrips razanii* | 1430 | 6.75 | Masumoto & Okajima, 2013 |
| *Lenkothrips guaraniticus* | 1200 | 7.31 | Cavalleri & Mound, 2014 |
| *Stenchaetothrips langkawiensis* | 1150 | 8.75 | Ng & Mound, 2012 |

### SUPPLEMENTARY MOVIES

**Movie 1.** Velocity vectors overlaid on out-of-plane vorticity ($\omega_z$) contours of bristled wing pairs during clap and fling, comparing the effect of increasing bristle diameter (*D*) from 0.1 mm to 0.3 mm. 10 equally spaced time instances are shown from start to end of clap, followed by 8 equally spaced time instances during fling.

**Movie 2.** Velocity vectors overlaid on out-of-plane vorticity ($\omega_z$) contours of bristled wing pairs during clap and fling, comparing the effect of increasing inter-bristle gap (*G*) from 1 mm to 2.1 mm. 10 equally spaced time instances are shown from start to end of clap, followed by 8 equally spaced time instances during fling.

**Movie 3.** Velocity vectors overlaid on out-of-plane vorticity ($\omega_z$) contours of bristled wing pairs during clap and fling, comparing the effect of increasing wing span (*S*) from 67.5 mm to 81 mm. 10 equally spaced time instances are shown from start to end of clap, followed by 8 equally spaced time instances during fling.
